# The Immune-Mutation Axis of Cancer Incidence: Insights from 30,000 TCR*β* Repertoires

**DOI:** 10.1101/2024.10.21.619470

**Authors:** H. Jabran Zahid, Matthew E. Lee, Ruth Taniguchi, Gheath Alatrash, Thomas McFall, Marco Garcia Noceda, Harlan Robins, Julia Greissl

## Abstract

Immunoediting posits that mutation and immunity jointly shape cancer evolution, yet their population-level interplay remains uncertain. Here we analyze T cell receptor (TCR) *β* repertoires from 30,000 individuals and find that TCR diversity, essential for recognizing and eliminating malignant cells, declines with age. This immune decline occurs 11 years later in females and coincides with their lower cancer incidence, suggesting a biological connection. To test this link, we formalize immunoediting as a quantitative model of carcinogenesis, relating the measured age-associated decline in TCR diversity to rising cancer incidence. We find that both mutational and immune processes shape cancer risk, with lower incidence in females attributable to delayed immune decline. Extending this analysis across subtypes uncovers structured patterns in cancer incidence that reflect the relative contributions of these processes. Cancers cluster along an emergent immune-mutation axis that aligns with known features of cancer biology and indicates convergent evolutionary dynamics. Together, our results establish a quantitative, population-level framework for immunoediting that connects direct measurements of immune competence to cancer risk, integrating the molecular mechanisms, evolutionary dynamics and incidence patterns of cancer to reveal a fundamental balance between mutation and immunity that underlies carcinogenesis.

## Introduction

The human population is rapidly aging, with the number of people over 60 years old expected to double by 2050 (*1*). A commensurate increase in cancer incidence and mortality is expected due to this demographic shift (*2*). As need for cancer prevention and treatment grows, understanding what drives cancer risk and why it rises so dramatically with age becomes increasingly urgent^1^. The idea that the immune system can prevent cancer dates back over a century (*4, 5*). The concept of cancer immunoediting builds on this foundation, describing how immune elimination of cells bearing visible neoantigens leads to the emergence of immune-evasive variants (*6*–*9*). This dynamic creates tension between mutational processes that drive cancer initiation and immune surveillance mechanisms that suppress it, though many aspects of this interaction remain poorly understood. To explore this interplay, we quantify immune decline and relate it to increasing cancer incidence with age, providing an integrated population-level perspective of immunoediting.

Quantifying immune competence is essential for understanding the role of the immune system in cancer. Alpha-beta T cells are the most abundant subtype of T cells, recognizing non-self-derived peptides through T cell receptor (TCR)-mediated interactions (*10, 11*). Consequently, a diverse repertoire of T cells confers immune competence, enabling responses to a broad range of immunological threats, including recognition and elimination of malignant cells (*12*–*21*). High repertoire diversity arises from random somatic rearrangements of TCR gene segments in the thymus, predominantly during childhood and adolescence before thymic involution (*22, 23*). Thus, TCR diversity provides a measure of the immune system’s functional capacity, which varies with age and other biological factors.

Aging impacts multiple components of the immune system and immunosenescence, broadly defined as the age-associated decline of immune competence, is characterized by a range of effects (*24*–*29*), including loss of TCR diversity (*30*–*34*). In elderly individuals, immune decline is believed to significantly contribute to higher disease incidence, poor health outcomes and inferior vaccine response (*35*–*41*). Loss of TCR diversity is a defining feature of immunosenescence and quantifying its decline with age provides a robust measure of diminishing immune competence.

Sex is a genetically defined biological factor that influences immune response and health outcomes (*42*). Males are more susceptible than females to many infectious diseases (*43, 44*) and they have poorer responses to vaccines (*45*). They also exhibit higher incidence rates for many cancers and worse survival outcomes (*46, 47*). While studies report that T cell repertoires decline more rapidly in males than in females (*48, 49*), significant uncertainties remain due to small sample sizes. Thus, sex-linked differences in immune decline, perhaps due to some combination of genetics, hormones and differences in thymic output (*42, 50*–*52*), may explain observed sex differences in cancer incidence. However, the relationship between sex, immune decline and cancer incidence remains poorly characterized partly because large-scale sex-stratified measurements of immune decline have been lacking. To address this gap, we quantify how immune decline varies with age and sex using TCR diversity measured in more than 30,000 subjects.

Mathematical models of carcinogenesis aim to describe the key processes that drive cancer onset. An early influential model by Armitage and Doll proposed that cancer arises from a series of discrete cellular mutations (*53*). Later models incorporated additional molecular mechanisms (*54*). However, immune decline and sex differences in incidence are rarely modeled explicitly. Palmer et al. (*55*) addressed this by modeling cancer incidence with an exponentially declining immune function tied to the timescale of thymic atrophy. They found that immune decline may be the dominant factor driving the increase in cancer risk with age, attributing sex differences in cancer incidence to sex-linked variation in thymic output. We extend their approach by directly quantifying immune decline using measurements of TCR diversity from deep repertoire sequencing, with sex providing a natural dimension of variation. By modeling how the forces of mutation and immunity jointly shape cancer incidence, our framework provides a mathematical formulation of cancer immunoediting constrained by population-level measurements.

T cell repertoire sequencing is a powerful probe of the adaptive immune system (*56*–*58*). Modern high-throughput methods allow sequencing of more than a million TCRs from a single blood draw (*59, 60*). While this captures only a small fraction of the estimated 5 *×* 10^11^ T cells in the human body (*61*), it provides an unbiased window into systemic immune changes across individuals. Leveraging large-scale measurements from over 30,000 subjects, we link immune decline and mutational processes to cancer incidence. This perspective refines our understanding of cancer immunoediting, with potential relevance for personalized medicines, effective immunotherapies and vaccines, and other strategies to address declining health with age (*62*–*65*).

## Results

We perform a cross-sectional analysis of a large homogeneously processed cohort of 30,430 unique subjects, approximately 20 to 80 years old, with TCR*β* repertoires sequenced as part of the T-Detect COVID test. This test was granted Emergency Use Authorization by the Food and Drug Administration^2^ for identifying the T cell immune response to SARS-CoV-2. We statistically analyze repertoires as a function of age and sex; the sequencing protocol is independent of these two variables. Because human leukocyte antigen (HLA) variation has minimal impact on overall TCR diversity (*66*), HLA genotype is not considered in this analysis. We interpret the total number of T cells sequenced, *S*, and the number of unique clonotypes (i.e., richness), *D*, as relative estimates of total repertoire size and total TCR diversity, respectively. These quantities are log-normally distributed and base 10 logarithms (referred to simply as “log”) are used throughout the analysis. Additionally, we define average clonality as the mean number of productive T cells per unique TCR rearrangement within a repertoire, given by the ratio of repertoire size to TCR diversity (i.e., *S/D*).

### Age and Sex Shape the T cell Repertoire

Figures 1A and 1B show age-associated trends in both repertoire size and TCR diversity, respectively. Between ages 20 and 80, repertoire size declines by a factor of 1.3 (0.1 dex^3^) and TCR diversity declines by a factor of 1.8 (0.25 dex). While decreasing repertoire size leads to loss of TCR diversity, greater loss is attributable to increasing average clonality of T cells (a factor of 1.4 or 0.15 dex; Figure 1C). After accounting for measurement uncertainty, we estimate that the peak-to-peak intrinsic biological scatter in TCR diversity for the central 90% of subjects increases by a factor of 2 to 5 (0.3 dex to 0.7 dex) between ages 20 and 80, respectively. Notably, at any given age, variation in TCR diversity across individuals is greater than the systematic change with age, particularly in older individuals.

**Figure 1.**
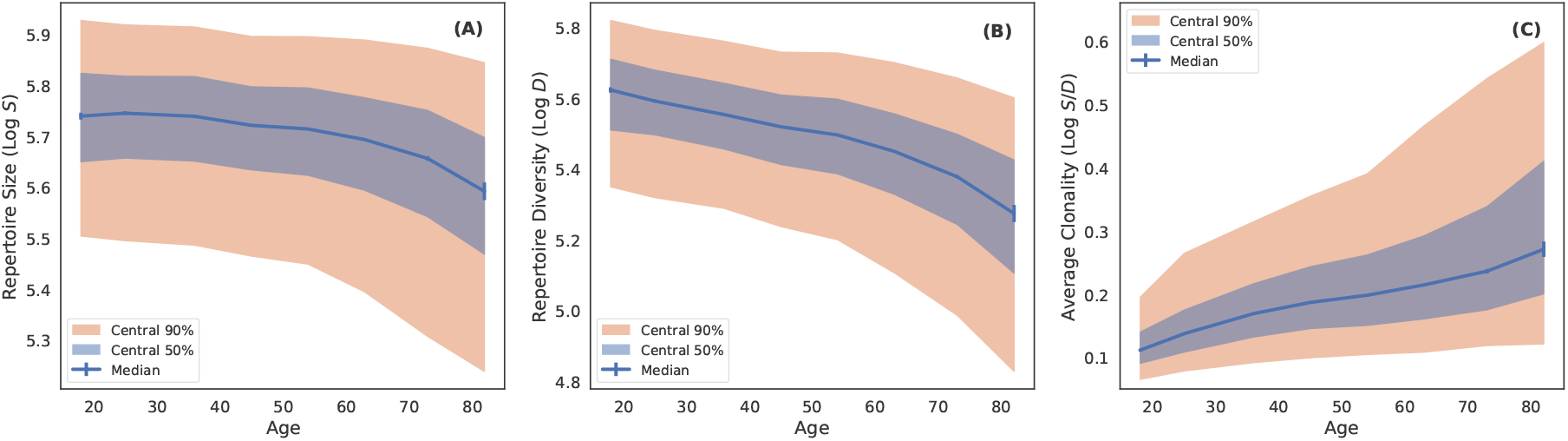
Repertoire size, TCR diversity and average clonality as a function of age. (A) Log of the total number of productive TCRs sequenced (Log *S*) in each repertoire as a function of age. The blue curve is the median in decade wide age bins. Error bars are bootstrapped and blue and orange shaded regions indicate the distribution of the central 50 and 90% of the data, respectively. (B) Log of TCR diversity (Log *D*) as a function of age. The binning procedure and shaded regions are the same as in (A). (C) Log of the average clonality of T cells in a repertoire as a function of age. For a given repertoire, average clonality is given by the ratio of repertoire size to TCR diversity (Log *S/D*). The binning procedure and shaded regions are the same as in (A). Binned data and errors are provided in Table 1.

Repertoire size and TCR diversity also vary systematically by sex (Figures 2A and 2B). Females younger than 45 years old have similar repertoire size and slightly higher TCR diversity than males. At older ages, females exhibit a slower decline in both measures. Similar age- and sex-associated trends in T cell fractions have been reported (*67, 68*) and we observe corresponding patterns in a smaller, independent cohort (see Supplementary Material Figure S2), reinforcing the robustness of these findings. Figures 2C and 2D show that the repertoire size and TCR diversity of females are both comparable to those of males who are 11.4 *±* 0.9 (1*σ*) years younger, demonstrating a delayed immune decline in females. This offset reflects a best-fit horizontal shift in adults and should not be taken as a uniform delay at all ages, since sex-based differences emerge primarily in middle age. While the origin of this trend remains uncertain, it is evident not only in the median values from tens of thousands of subjects but also in the consistent scatter between sexes, suggesting that observed differences likely reflect a mechanism that generally affects the male and/or female population, such as more enduring naive T cell production in females (*50*).

**Table 1:** Repertoire size and TCR diversity as a function of age. The first and second columns give the number of subjects in the bin and median age, respectively. The third and fifth columns give the median log of repertoire size, *S*, and log TCR diversity, *D*, respectively. The fourth and sixth columns give the errors on the median log repertoire size and log TCR diversity, respectively. Errors are 1*σ* uncertainties determined from 10,001 bootstrapped realizations. Measurements for all subjects, male subjects and female subjects are given in the table in descending order.

| N Subjects | Median Age | $\text{Log}_{10}S$ | Error $\text{Log}_{10}S$ | $\text{Log}_{10}D$ | Error $\text{Log}_{10}D$ |
| --- | --- | --- | --- | --- | --- |
| All Subjects |  |  |  |  |  |
| 561 | 18 | 5.739 | 0.007 | 5.623 | 0.008 |
| 1591 | 25 | 5.745 | 0.004 | 5.592 | 0.005 |
| 5043 | 36 | 5.739 | 0.002 | 5.553 | 0.002 |
| 7608 | 45 | 5.721 | 0.002 | 5.519 | 0.002 |
| 8252 | 54 | 5.714 | 0.002 | 5.496 | 0.002 |
| 5491 | 63 | 5.693 | 0.002 | 5.449 | 0.003 |
| 1647 | 73 | 5.656 | 0.004 | 5.377 | 0.005 |
| 237 | 82 | 5.592 | 0.017 | 5.274 | 0.023 |
| Male Subjects |  |  |  |  |  |
| 272 | 18 | 5.736 | 0.009 | 5.617 | 0.012 |
| 754 | 25 | 5.735 | 0.006 | 5.576 | 0.007 |
| 2564 | 36 | 5.739 | 0.003 | 5.542 | 0.003 |
| 3642 | 45 | 5.722 | 0.002 | 5.507 | 0.003 |
| 3777 | 54 | 5.694 | 0.002 | 5.460 | 0.003 |
| 2475 | 63 | 5.664 | 0.003 | 5.408 | 0.005 |
| 779 | 73 | 5.624 | 0.010 | 5.328 | 0.008 |
| 108 | 82 | 5.542 | 0.020 | 5.170 | 0.030 |
| Female Subjects |  |  |  |  |  |
| 288 | 18 | 5.742 | 0.010 | 5.625 | 0.009 |
| 833 | 25 | 5.752 | 0.005 | 5.605 | 0.007 |
| 2461 | 36 | 5.738 | 0.003 | 5.567 | 0.004 |
| 3950 | 45 | 5.721 | 0.003 | 5.532 | 0.003 |
| 4448 | 54 | 5.731 | 0.002 | 5.525 | 0.003 |
| 3010 | 63 | 5.715 | 0.003 | 5.483 | 0.003 |
| 866 | 72 | 5.683 | 0.006 | 5.417 | 0.009 |
| 128 | 82 | 5.647 | 0.017 | 5.329 | 0.030 |

**Figure 2.**
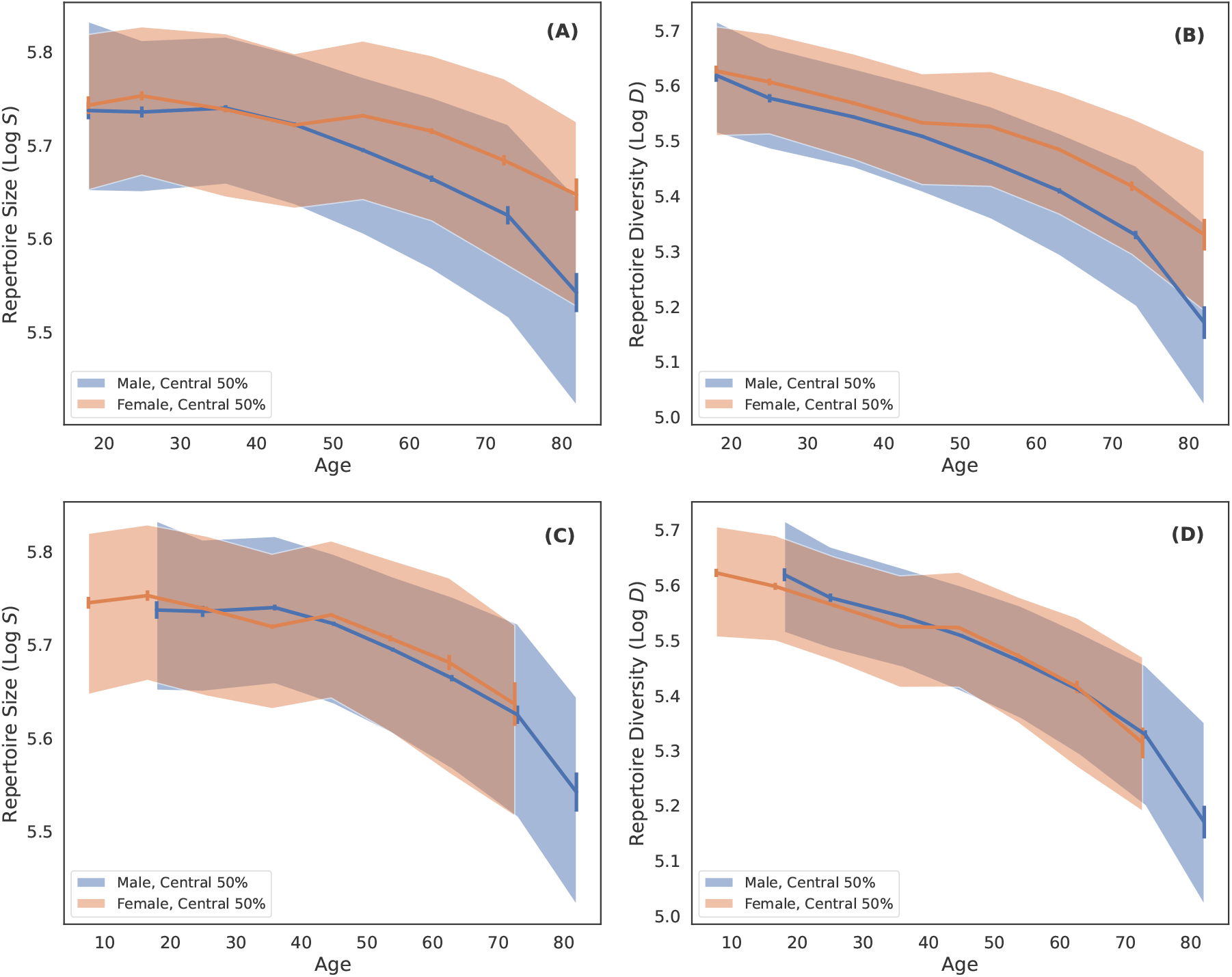
Repertoire size and TCR diversity as a function of age and sex. (A) Log of the total number of productive TCRs sequenced (Log *S*) as a function of age and sex. Blue and orange curves are the median in decade wide age bins for males and females, respectively. Error bars are bootstrapped and shaded regions indicate the distribution of the central 50% of the data points. (B) Log of the TCR diversity as a function of age (Log *D*) for male (blue) and female (orange) subjects. The binning procedure and shaded regions are the same as in (A). (C) and (D) are the same as (A) and (B), respectively, but with female samples shifted to younger ages by 11.4 years and rebinned for display. Binned data and errors are provided in Table 1.

### A Model Linking Immune Decline, Sex and Cancer Incidence

Using cancer incidence data from the Surveillance, Epidemiology and End Results (SEER) database together with TCR diversity measured from our cohort, we model cancer incidence as a function of age, sex and TCR diversity. Cancer incidence and TCR diversity show opposing trends such that incidence increases with age and is lower in females, whereas TCR diversity declines with age and is higher in females (compare Figures 2B and 3A). Given the central role of T cells in eliminating cancerous cells, we hypothesize that loss of TCR diversity may contribute to rise in cancer incidence with age (*55, 69*–*72*). If so, then delayed immune decline in females may help explain their lower incidence of cancer.

To link immune decline, sex and cancer incidence, we develop a biologically grounded correlative model of carcinogenesis designed for simplicity and interpretability. Here we define carcinogenesis as the initial failure of immune clearance, corresponding to the elimination phase of the cancer immunoediting framework, when a transformed cell evades detection (*6, 7*). While our model is scoped to this early phase, TCR diversity likely influences later stages of tumor evolution as well, including equilibrium and escape. To the extent that these later phases depend on immune competence, our framework may also capture relevant signals of those processes.

We mathematically model two key biological processes contributing to carcinogenesis with age: (1) accumulation of driver mutations and (2) diminishing immune surveillance. The mutational component follows the Armitage and Doll framework, a simple power-law function of age in which the exponent is interpreted as quantifying the number of discrete driver mutations required for carcinogenesis (*53*). The immune component is modeled as an exponential function of TCR diversity (*73, 74*). To understand this dependence, let *p* denote the probability that a random TCR recognizes a given antigen. Then the probability that an antigenic mutation is not recognized by 𝒟 unique TCRs is 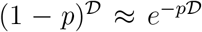 for small *p*, a reasonable assumption given the rarity of TCR-antigen matches. Thus, the likelihood of immune elimination failure increases exponentially as TCR diversity declines, a functional dependence that emerges naturally from the probabilistic basis of TCR-antigen recognition. To reproduce the plateau or decline in incidence at advanced ages (*75, 76*), we include a logistic suppression term that improves goodness-of-fit with minimal covariation with other parameters. Our model is a product of these factors:

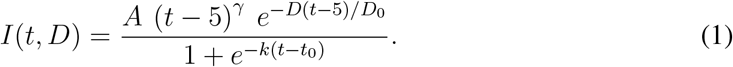

Here *I*(*t, D*) is cancer incidence as a function of age *t* and measured TCR diversity *D*, and *A, γ, D*_0_, *k* and *t*_0_ are free parameters. Normalizing *D* by *D*_0_ means our TCR diversity measurements require only relative accuracy, which our data provide. The 5-year age offset accounts for latency between the elimination phase of immunoediting, which our model represents, and the escape phase, which corresponds to clinical detection and is what our model fits. In other words, cancer incidence observed at a given age reflects mutation burden and TCR diversity at an earlier time, prior to clinical detection. Estimates from screening cohorts, registry analyses and genomic reconstructions suggest that this preclinical period varies widely by cancer type, ranging from only a few years in rapidly progressing tumors to one or two decades in more indolent cancers (*77*–*80*). Larger values for this latency parameter effectively shift explanatory power from the mutational component to the immune component in our model. Although we would ideally treat it as a free parameter, it cannot be reliably constrained because of parameter covariance (see Materials & Methods). Our primary conclusions are robust to reasonable variations in this parameter, so to remain conservative in estimating the immune contribution, we fix the latency at 5 years.

We model cancer incidence as a function of age and sex, aggregating incidence across subtypes. We incorporate TCR diversity, represented by *D*, into the model (Equation 1), using the median age-dependent diversity curves measured separately for males and females (Figure 2B) to fit the corresponding male and female incidence data. This simple approach yields results consistent with individual-level modeling, where risk is computed per subject from age and TCR diversity and then aggregated by age and sex (see Materials & Methods). Importantly, sex is not an explicit input variable of the model; a single set of parameters is optimized to fit both male and female incidence curves. All sex differences in the fit arise solely from the sex dependence of TCR diversity, which is the only sex-specific input. In other words, our model quantitatively maps sex differences in TCR diversity to sex differences in cancer incidence. This is notable because both the accumulation of mutations and declining TCR diversity contribute to rising cancer incidence with age, but only TCR diversity varies with sex. This sex dependence breaks the degeneracy between components, allowing us to independently isolate risk attributable to immune decline and to accumulation of mutations. Model parameters are constrained using non-linear least squares optimization, yielding best-fit values with 95% confidence intervals (Figure 3A, Table 2).

**Table 2:** Parameter fits by cancer subtype. A values are always shown in scientific notation with two significant digits. D_0_ values are shown in scientific notation with three significant digits. Error bars are scaled to the same exponent as the central value. *γ* has two decimals, *k* two significant digits (or -1 if not fit), and *t*_0_ one decimal.

| Cancer Subtype | A | $\gamma$ | $D_0$ | $k$ | $t_0$ |
| --- | --- | --- | --- | --- | --- |
| Aggregated Cancers (Initial Selection) | $2.5^{+3.7}_{-1.8} \times 10^0$ | $2.22^{+0.22}_{-0.15}$ | $8.99^{+1.3}_{-0.9} \times 10^4$ | $-0.20^{+0.03}_{-0.03}$ | $82.3^{+1.7}_{-1.7}$ |
| Acute Myeloid Leukemia | $5.6^{+12.5}_{-4.6} \times 10^{-1}$ | $1.45^{+0.33}_{-0.22}$ | $8.85^{+1.48}_{-0.80} \times 10^4$ | -1 | $84.2^{+3.5}_{-0.6}$ |
| Chronic Lymphocytic Leukemia | $7.6^{+41.5}_{-6.8} \times 10^{-7}$ | $5.28^{+0.53}_{-0.40}$ | $7.30^{+1.07}_{-0.85} \times 10^4$ | $-0.13^{+0.01}_{-0.02}$ | $66.0^{+3.3}_{-4.1}$ |
| Chronic Myeloid Leukemia | $1.1^{+0.8}_{-0.8} \times 10^1$ | $0.42^{+0.22}_{-0.09}$ | $9.82^{+1.51}_{-0.68} \times 10^4$ | $-0.25^{+0.11}_{-0.75}$ | $87.6^{+4.6}_{-3.4}$ |
| Colorectal Adenocarcinoma MSS | $1.4^{+3.4}_{-1.0} \times 10^{-4}$ | $4.06^{+0.37}_{-0.32}$ | $1.28^{+0.20}_{-0.16} \times 10^5$ | $-0.067^{+0.011}_{-0.015}$ | $62.8^{+8.4}_{-12.2}$ |
| Diffuse Large B Cell Lymphoma | $1.3^{+2.6}_{-1.0} \times 10^{-1}$ | $1.88^{+0.26}_{-0.19}$ | $1.04^{+0.17}_{-0.12} \times 10^5$ | $-0.56^{+0.24}_{-0.44}$ | $84.5^{+1.3}_{-1.0}$ |
| Follicular Lymphoma | $1.8^{+0.7}_{-0.6} \times 10^{-5}$ | $3.57^{+0.09}_{-0.08}$ | $2.91^{+0.35}_{-0.22} \times 10^5$ | $-0.092^{+0.008}_{-0.010}$ | $75.5^{+1.7}_{-1.7}$ |
| Gastric Adenocarcinoma | $2.4^{+10.5}_{-2.1} \times 10^0$ | $1.81^{+0.40}_{-0.27}$ | $5.35^{+0.78}_{-0.63} \times 10^4$ | $-0.15^{+0.04}_{-0.04}$ | $76.9^{+3.0}_{-4.2}$ |
| Kidney Chromophobe Carcinoma | $2.2^{+2.8}_{-1.5} \times 10^{-2}$ | $1.93^{+0.25}_{-0.17}$ | $9.52^{+1.19}_{-0.72} \times 10^4$ | $-0.16^{+0.02}_{-0.03}$ | $73.8^{+1.7}_{-1.8}$ |
| Kidney Clear Cell Carcinoma | $2.0^{+5.0}_{-1.6} \times 10^{-1}$ | $2.47^{+0.31}_{-0.23}$ | $6.54^{+0.81}_{-0.59} \times 10^4$ | $-0.15^{+0.02}_{-0.02}$ | $69.4^{+2.2}_{-2.3}$ |
| Kidney Papillary Carcinoma | $4.3^{+50.1}_{-4.1} \times 10^2$ | $1.16^{+0.61}_{-0.44}$ | $3.32^{+0.36}_{-0.33} \times 10^4$ | $-0.23^{+0.03}_{-0.05}$ | $68.3^{+2.1}_{-2.6}$ |
| Lung Adenocarcinoma | $8.1^{+0.2}_{-0.7} \times 10^{-10}$ | $6.45^{+0.02}_{-0.01}$ | $3.59^{+0.22}_{-0.16} \times 10^5$ | $-0.14^{+0.01}_{-0.01}$ | $72.8^{+0.4}_{-0.4}$ |
| Lung Small Cell Carcinoma | $2.3^{+81.2}_{-0.9} \times 10^{-12}$ | $7.75^{+0.12}_{-0.88}$ | $2.88^{+0.36}_{-0.64} \times 10^5$ | $-0.16^{+0.01}_{-0.02}$ | $67.1^{+3.8}_{-0.6}$ |
| Lung Squamous Cell Carcinoma | $1.6^{+8.2}_{-1.4} \times 10^{-8}$ | $6.45^{+0.55}_{-0.35}$ | $6.54^{+0.99}_{-0.86} \times 10^4$ | $-0.20^{+0.02}_{-0.02}$ | $70.8^{+1.5}_{-2.2}$ |
| Malignant Gliomas | $9.8^{+14.0}_{-7.1} \times 10^{-1}$ | $1.38^{+0.22}_{-0.14}$ | $9.17^{+1.42}_{-0.91} \times 10^4$ | $-0.23^{+0.02}_{-0.03}$ | $80.6^{+1.2}_{-1.2}$ |
| Melanoma | $8.9^{+15.8}_{-7.2} \times 10^1$ | $0.96^{+0.33}_{-0.19}$ | $6.71^{+0.87}_{-0.46} \times 10^4$ | $-0.15^{+0.03}_{-0.04}$ | $79.3^{+2.7}_{-2.6}$ |
| Multiple Myeloma | $8.5^{+13.4}_{-5.9} \times 10^{-7}$ | $4.87^{+0.28}_{-0.20}$ | $1.18^{+0.16}_{-0.12} \times 10^5$ | $-0.11^{+0.01}_{-0.02}$ | $70.3^{+2.7}_{-3.5}$ |
| Pancreatic Adenocarcinoma | $2.4^{+2.9}_{-1.4} \times 10^{-9}$ | $6.20^{+0.21}_{-0.16}$ | $1.78^{+0.30}_{-0.23} \times 10^5$ | $-0.12^{+0.01}_{-0.01}$ | $69.0^{+1.7}_{-1.9}$ |
| Soft Tissue Sarcomas | $4.5^{+3.5}_{-3.0} \times 10^0$ | $0.73^{+0.19}_{-0.10}$ | $9.31^{+1.24}_{-0.64} \times 10^4$ | $-0.28^{+0.08}_{-0.24}$ | $86.1^{+2.2}_{-1.5}$ |

**Figure 3.**
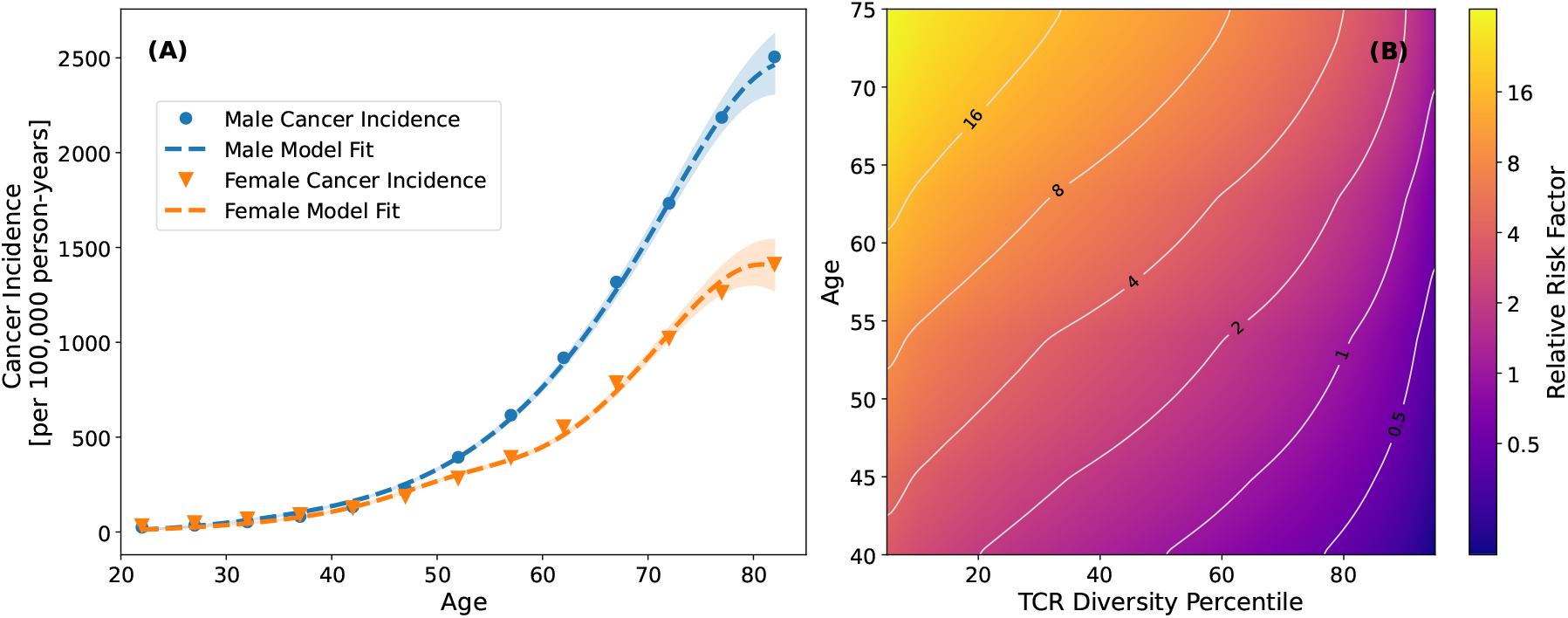
Modeling cancer incidence as a function of age and TCR diversity. (A) Blue and orange points are cancer incidence (per 100,000 person-years) for males and females, respectively. Cancer incidence is modeled as a function of age and TCR diversity using Equation 1 and the median TCR diversity measured for each sex (see Figure 2B). Dotted lines are the best-fit model and shaded regions are 95% confidence intervals. (B) A heatmap showing relative cancer risk as a function of age and TCR diversity percentile. TCR diversity percentiles indicate the fraction of subjects at a given age who have TCR diversity values below that specific value. Relative cancer risk is calculated as the cancer incidence derived using our best-fit model normalized to cancer incidence of a 40 year old with median TCR diversity. Contours of constant relative risk are shown in gray.

We factor cancer risk associated with accumulation of mutations and immune decline by noting that cancer incidence fit by a single power-law scales as *t*^4.6^, cf. Palmer et al. (*55*). In our model, risk associated with accumulation of mutations scales as *t*^2.22^ and immune decline accounts for the remainder of the risk. This decomposition is broadly consistent with Palmer et al. and supports the view that both processes contribute significantly to age-related cancer risk. We partition cancer risk into two components: (1) risk attributable to aging, reflecting contributions from both the accumulation of mutations and systematic immune decline and (2) risk arising from variation in TCR diversity among individuals of the same age (see Materials & Methods). Figure 3B shows our partitioning of relative cancer risk as a function of age and TCR diversity percentile. At the median TCR diversity (Figure 1B), a 75-year-old faces a ∼12-fold higher risk of cancer than a 40-year-old, with this increase driven equally by the accumulation of mutations and loss of TCR diversity. Conversely, a 75-year-old in the tenth percentile has a factor of ∼3 lower TCR diversity than a peer in the ninetieth percentile, resulting in a ∼15-fold higher risk of cancer solely due to this difference. This disparity underscores the exponential dependence of cancer risk on TCR diversity. If individuals were able to maintain the median TCR diversity observed at age 40 into old age (e.g., through medical interventions), our model predicts a four-fold reduction in cancer incidence at age 75. By contrast, in the hypothetical absence of all T cells (*D* = 0), our model predicts cancer incidence would be significantly higher than what is observed in the general population. While no individual completely lacks T cells, studies report that immunocompromised or thymectomized subjects have higher risk of cancer (*81*–*83*).

We show that immune decline may contribute significantly to cancer risk. Measurements of TCR diversity could provide a practical means of identifying individuals at elevated risk, for whom preventative medical interventions may be warranted. The risk quantification presented in Figure 3B demonstrates how such individuals can be identified in a straightforward manner.

### An Immune-Mutation Axis Organizes Cancer Incidence

We evaluate the biological relevance of our model by testing whether variation in its components reflects known differences across cancer subtypes. Our aim is to validate the model and contextualize its interpretation, yielding a population-level perspective on cancer biology grounded in the combination of large-scale immunosequencing and epidemiological data.

Immune decline is likely a general risk factor for cancer. To test this hypothesis, we examine whether females show lower incidence across subtypes, consistent with delayed immune decline. Because sex differences can also arise from anatomy, hormones, behavior or sex-biased viral exposures, we restrict our analysis to cancers with minimal potential confounding (see Materials & Methods). Of the 18 subtypes included, 15 strictly meet these criteria; the remaining three—lung adenocarcinoma, lung squamous cell carcinoma and small cell lung carcinoma— are retained with caution due to smoking-related confounding, which has historically been more prevalent in males (*84*–*86*). In these cancers, the model may misattribute behavioral differences to immune decline, overestimating immune sensitivity and underestimating mutational contributions. We nonetheless include them because their cautious interpretation strengthens the generality of our conclusions.

We fit age-specific cancer incidence for males and females across 18 subtypes using the same model and procedure described previously (Table 2). Under the hypothesis that immune decline generally contributes to cancer risk, subtypes selected to minimize sex confounders are expected to show higher incidence in males. Consistent with this expectation, all 18 subtypes do (Figure 4). Supporting this pattern, a large cohort study of thymectomized adults reported a two-fold increase in cancer incidence compared to matched surgical controls (*83*). Although that study lacked resolution of individual cancer subtypes, several large cohort studies of HIV infection, solid organ transplantation and primary immunodeficiencies report elevated incidence across nearly all of the subtypes we analyze (*87*–*90*). Notably, in immunosuppressed transplant recipients the usual male excess in incidence is reduced or absent, consistent with immunosuppression equalizing immune capacity between sexes (*91*). Taken together, these observations are consistent with our model’s prediction that immune decline contributes to sex differences in cancer incidence, although confounding factors and inconsistent histologic resolution in studies of immunosuppressed individuals preclude detailed quantitative comparison.

**Figure 4.**
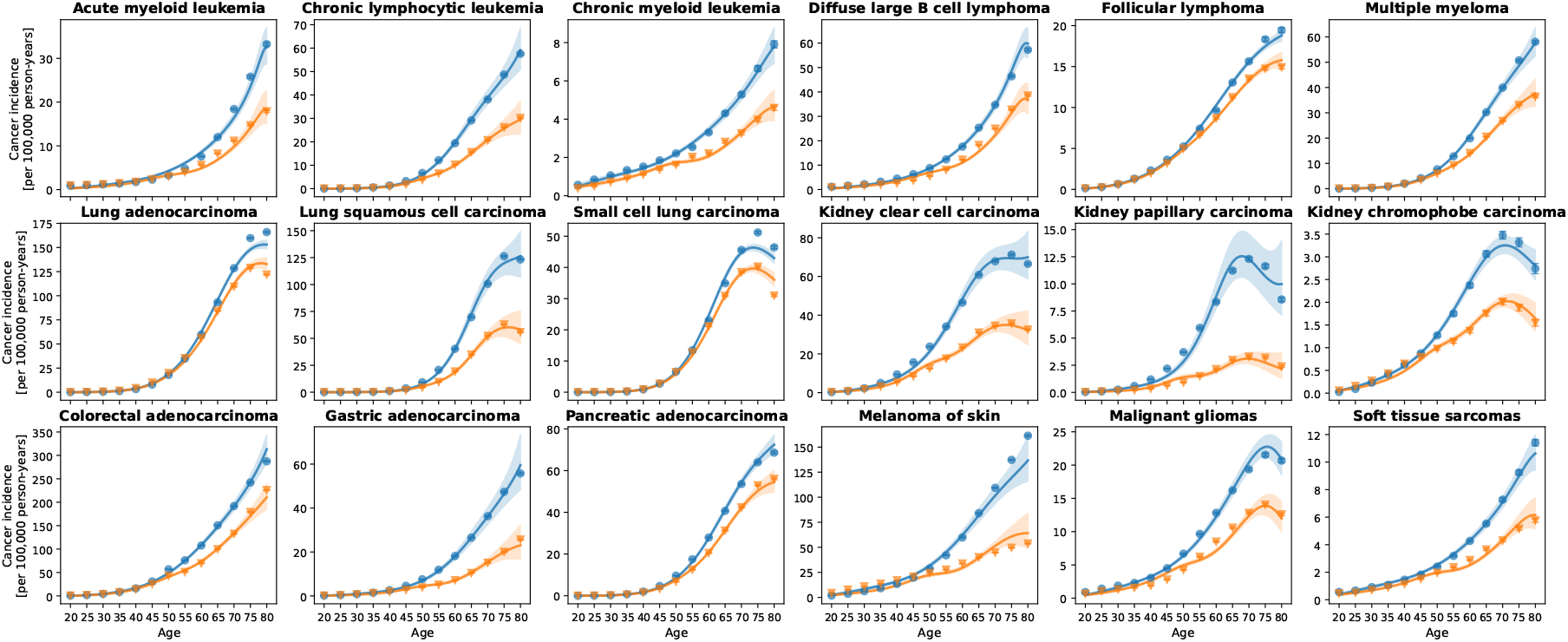
Model fits to cancer incidence data across 18 subtypes. The data points show age-dependent incidence in males (blue circles) and females (orange triangles). Blue and orange curves are fit to the male and female incidence data, with a shared parameter set fit for each cancer subtype using observed sex-specific TCR diversity. Shaded regions indicate 95% confidence intervals. Cancers are broadly grouped by organ system.

To assess model validity more directly, we examine whether the Armitage and Doll (*53*) interpretation of the power-law term, that the exponent corresponds to the number of required driver mutations, applies in our framework. Across cancer subtypes, our fit of *γ* ranges from 0.4 to 7.8. In comparison, Martincorena et al. (*92*) estimated driver mutation counts from tumor genetic sequencing using dN/dS ratios and other evolutionary signatures. They find that carcinogenesis is driven by a small number of driver mutations in known cancer genes per tumor, with counts systematically varying across subtypes. Their estimates are directly comparable to our values of *γ* and across nine overlapping subtypes we observe a significant positive correlation (Spearman *ρ* = 0.7, *p* = 0.04; see Methods). Although statistical power is limited, this agreement provides independent molecular validation of our interpretation.

To further aid interpretation and validation, we visualize how fit parameters map to biological processes. For each subtype, we partition the increase in cancer risk between ages 40 and 70 into components attributable to the accumulation of mutations (*M*_*c*_) and immune decline (*I*_*c*_), assuming median TCR diversity at each age for the latter. Figure 5 reveals a natural division between subtypes that are *mutation-dominated*, where incidence is strongly shaped by accumulating mutations, and those that are *immune-modulated*, where immune engagement plays a larger role. The clustered distribution of cancers in incidence space defines an *immune-mutation axis*, reflecting how the balance between these forces shifts across tissues and subtypes. Notably, *M*_*c*_ varies more across subtypes than *I*_*c*_ and overall incidence (summed over age and sex) correlates with *M*_*c*_ and *γ* (Spearman *ρ* = 0.6, *p* = 0.01; Table 3), but not with *I*_*c*_. These results suggest that subtype differences are influenced more strongly by the mutational component, though the immune component remains essential. The correlation of overall incidence with *γ* may reflect that tissues with higher cellular turnover generate greater mutational capacity (*93, 94*), increasing both the probability of malignant transformation and the feasibility of multi-driver trajectories. While the biology forms a continuum, grouping subtypes by the dominant factors shaping incidence provides a framework for interpreting incidence patterns in the context of broader cancer biology.

**Table 3:** Age 40 to 70 risk factors by cancer subtype and summed incidence. The first column gives the cancer subtype and columns two, three and four give the overall fold-change in increased risk at age 70 compared to age 40, risk associated with immune decline (*I*_*c*_) and risk associated with accumulation of mutation (*M*_*c*_), respectively. Column five gives the ratio of mutational-to-immune risk ratio. Column six gives the aggregated overall incidence (per 100,000 person-years) summed across both sexes for the age range we evaluate. Values are sorted in descending order of immune-to-mutational risk ratio.

| Cancer Subtype | $F_r$ | $I_c$ | $M_c$ | $M_c/I_c$ | $\Sigma$ Incidence | Category |
| --- | --- | --- | --- | --- | --- | --- |
| Small Cell Lung Carcinoma | 65.8 | 1.38 | 121 | 88.0 | 410 | Mutation-dominated |
| Lung Adenocarcinoma | 41.9 | 1.29 | 54.0 | 41.8 | 1245 | Mutation-dominated |
| Pancreatic Adenocarcinoma | 37.8 | 1.68 | 46.5 | 27.7 | 517 | Mutation-dominated |
| Lung Squamous Cell Carcinoma | 120 | 4.11 | 54.2 | 13.2 | 736 | Mutation-dominated |
| Multiple Myeloma | 23.4 | 2.18 | 20.4 | 9.35 | 379 | Mutation-dominated |
| Chronic Lymphocytic Leukemia | 36.1 | 3.55 | 26.3 | 7.41 | 333 | Mutation-dominated |
| Follicular Lymphoma | 8.13 | 1.37 | 9.13 | 6.64 | 179 | Mutation-dominated |
| Colorectal Adenocarcinoma MSS | 11.8 | 2.06 | 12.3 | 5.99 | 2028 | Mutation-dominated |
| Diffuse Large B Cell Lymphoma | 7.80 | 2.44 | 3.20 | 1.31 | 373 | Immune-modulated |
| Kidney Chromophobe Carcinoma | 5.62 | 2.64 | 3.30 | 1.25 | 33 | Immune-modulated |
| Kidney Clear Cell Carcinoma | 9.18 | 4.10 | 4.63 | 1.13 | 611 | Immune-modulated |
| Acute Myeloid Leukemia | 6.98 | 2.84 | 2.46 | 0.864 | 187 | Immune-modulated |
| Malignant Gliomas | 5.90 | 2.74 | 2.35 | 0.857 | 199 | Immune-modulated |
| Soft Tissue Sarcomas | 4.19 | 2.70 | 1.57 | 0.583 | 81 | Immune-modulated |
| Gastric Adenocarcinoma | 12.7 | 5.63 | 3.07 | 0.546 | 308 | Immune-modulated |
| Chronic Myeloid Leukemia | 3.27 | 2.56 | 1.29 | 0.505 | 65 | Immune-modulated |
| Melanoma | 5.73 | 3.97 | 1.81 | 0.456 | 1025 | Immune-modulated |
| Kidney Papillary Carcinoma | 13.5 | 16.2 | 2.05 | 0.127 | 84 | Immune-modulated |

**Figure 5.**
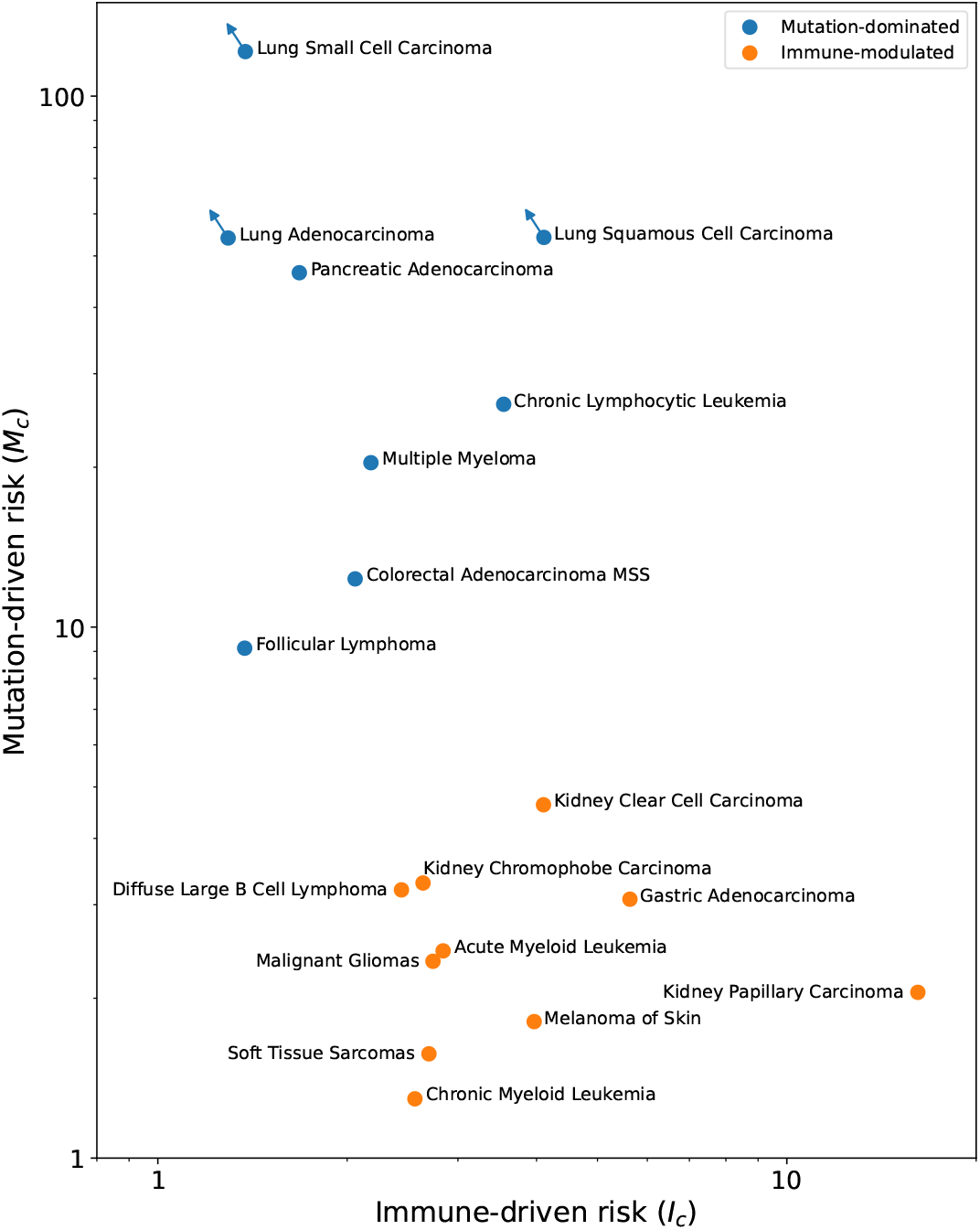
The immune-mutation axis organizes cancer subtypes by their dominant biological drivers. Each point represents a cancer subtype positioned by the model-inferred contributions of the accumulation of mutations (*M*_*c*_, x-axis) and immune decline (*I*_*c*_, y-axis) to the increase in cancer risk between ages 40 and 70. Values are given in Table 3 and are calculated using model parameters in Table 2. Cancers are colored into two groups that emerge from their separation on this plot. Lung cancers are shown with arrows to indicate possible adjustment for smoking confounding. The Cancers cluster along the immune-mutation axis, revealing a structured continuum of cancer behavior governed by the balance of mutation and immunity.

We evaluate the immune-mutation axis against independent measures of immune activity. Rooney et al. (*95*) quantified cytolytic activity in tumors relative to normal tissue using expression levels of perforin and granzyme A. Notably, their relative measure contrasts with the hot-cold tumor classification, which reflects absolute immune infiltration (*96*). Subtypes we identify as mutation-dominated, including colorectal adenocarcinoma, lung adenocarcinoma and lung squamous cell carcinoma, show lower cytolytic activity in tumors than in normal tissue. In contrast, immune-modulated cancers such as melanoma, kidney carcinomas, gliomas and gastric adenocarcinoma, show higher cytolytic activity in tumors. The complete concordance across overlapping subytpes demonstrates that the immune-mutation axis captures functional immune engagement.

Blood malignancies illustrate how cancers of shared hematopoietic origin diverge along the immune-mutation axis. Chronic lymphocytic leukemia, follicular lymphoma and multiple myeloma fall in the mutation-dominated region, consistent with stepwise accumulation of driver mutations and limited immune clearance (*97*–*99*). These cancers typically follow indolent courses. By contrast, acute myeloid leukemia, chronic myeloid leukemia and diffuse large B-cell lymphoma cluster in the immune-modulated region and show strong evidence of immune engagement, including escape mechanisms such as HLA downregulation and checkpoint ligand upregulation (*100*–*104*). Although immune escape occurs across all blood cancers, the tempo and nature of selective pressures align broadly with position on the axis. Similar patterns appear in solid tumors, indicating that the immune-mutation axis captures meaningful differences in evolutionary dynamics across subtypes, including those of related lineage and environment. Finally, we test whether the model captures finer variation within single sites. In the kidney, all subtypes fall in the immune-modulated region. Clear cell and chromophobe carcinomas sit nearer the mutation-dominated boundary, while papillary carcinoma lies further toward the immune-modulated end. This ordering is consistent with papillary carcinoma having the lowest mutational contribution, clear cell carcinoma showing a stronger mutational component alongside immune involvement and chromophobe carcinoma exhibiting the most mutation-driven profile (*105*–*107*). In the lung, all subtypes fall in the mutation-dominated region. Adenocarcinoma and squamous cell carcinoma lie nearer the immune-modulated boundary, while small cell carcinoma sits at the extreme mutation-dominated end. This placement matches known biology, with adenocarcinoma and squamous subtypes showing lower mutational burden and slower trajectories and small cell carcinoma exhibiting both high burden and rapid progression (*108, 109*). Smoking-related confounding may overestimate immune contributions in lung cancer, particularly squamous and small cell carcinoma, though this contribution is modest regardless and our interpretation remains robust. Thus, the immune-mutation axis resolves biologically meaningful heterogeneity even within a single site.

Mutation and immunity, the core elements of immunoediting, shape cancer risk across subtypes. By combining population-level TCR sequencing with epidemiological data, we derive a mathematical formulation of cancer immunoediting that defines an immune-mutation axis along which cancer subtypes cluster. Situating this axis within the context of molecular studies provides an integrated framework for interpreting evolutionary cancer biology across modalities.

## Discussion

Age and sex strongly influence cancer incidence, but the biological mechanisms driving this relationship remain poorly understood. To address this gap, we analyze ∼30,000 deeply sequenced T cell repertoires from individuals aged 20 to 80, showing that TCR diversity declines with age. This decline occurs about 11 years later in females than in males and coincides with their lower cancer incidence across diverse subtypes. Our analysis suggests that loss of immune competence contributes to age-related cancer risk and that sex differences in incidence arise from delayed immune decline in females. While compromised immunity is a well-established determinant of susceptibility to virally driven cancers (*81, 88, 110*), our findings suggest that this principle applies broadly across cancer subtypes. By modeling carcinogenesis, we provide a biologically grounded mathematical framework for understanding how mutation and immunity jointly shape cancer risk across diverse subtypes.

Several large-scale studies reinforce our results and provide complementary perspectives. Because immunosuppression weakens immune function thereby reducing immune differences between males and females, our model predicts higher overall cancer incidence and an attenuated difference between sexes in immunosupressed individuals, patterns borne out in large cohort studies (*83, 91*). Population-scale genomic analyses inferring T cell fractions from whole genome sequencing (WGS) (*111*) further support our findings. Poisner et al. (*67*) estimated T cell fractions in two large independent cohorts and found age- and sex-associated trends that closely mirror our measurements of repertoire size, a directly comparable metric. Using the same approach, the ImmuneLENS study found similar trends and reported that individuals with cancer exhibit significantly lower T cell fractions, with T cell abundance proving more prognostic of 5-year survival than tumor-infiltrating lymphocytes (*68*). Complementing these WGS studies, we use TCR diversity to quantify immune competence, which is a more direct and precise estimate of the immune system’s functional capacity.

Our analysis reveals that across subtypes, cancer incidence reflects the dynamics of immunoediting, captured quantitatively by position along the immune-mutation axis. Subtype positions align with known features of cancer biology, including driver burden, cytolytic activity, immune escape and selective pressures. Cancers naturally cluster along this axis, suggesting convergent evolutionary strategies. In tissues with greater capacity for driver accumulation, the probability of assembling multi-driver trajectories is higher, enabling mutation-dominated cancers to arise despite immune pressure. Such cancers often progress through canonical pathways and are associated with higher overall incidence perhaps reflecting greater mutational opportunity (*93*). In tissues with lower capacity, limited opportunities for driver accumulation make progression more mutationally constrained and thus more dependent on immune decline. This produces immune-modulated cancers with fewer drivers and progression more strongly shaped by immune engagement. The interplay between mutation and immunity is context dependent, as illustrated by the varying roles of canonical driver mutations. In pancreatic adenocarcinoma, KRAS mutations are nearly universal, primarily serving as initiating steps in multi-driver trajectories (*112*), consistent with the high driver accumulation that characterizes mutation-dominated cancers. By contrast, in immune-modulated cancers such as melanoma, BRAF mutations are also canonical drivers, yet progression remains strongly shaped by immune pressure; here, fewer driver mutations are required and abundant UV-induced mutations create strong immune engagement that makes surveillance and escape central to disease evolution (*95, 113*). The immune-mutation axis emerges as a population-level consequence of immunoediting, revealing how evolutionary dynamics shape cancer incidence across tissues and subtypes.

Oversimplification of the interplay between mutation and immunity may help explain why widely used classification schemes often perform inconsistently. Tumor mutational burden (TMB) was promoted as a pan-cancer predictor of response to immune checkpoint inhibition (*114*–*117*), which has been a therapeutic revolution in cancer care. However, large analyses show it fails in many cancers (*118, 119*). This is partly because TMB is an imperfect proxy for immune visibility. For instance, we identify kidney clear cell carcinoma and gliomas as immune-modulated, with immune engagement evident despite modest TMB (*95*), whereas lung small cell carcinoma and pancreatic adenocarcinoma are both mutation-dominated, showing little immune involvement and poor immunotherapy response despite very different TMBs (*108, 112, 118*). Moreover, despite their distinct functional roles, TMB combines driver and passenger mutations, with passengers vastly outnumbering drivers (*92*). Melanomas are immune-modulated despite very high TMB, with their evolution shaped by only a few recurrent drivers (*92*), consistent with their small *γ* value. Similarly, ambiguity may arise from erroneously conflating T cell infiltration with immune engagement. Microsatellite-stable colorectal adenocarcinoma often shows infiltration, yet its evolution is mutation-dominated and it remains resistant to checkpoint blockade (*120, 121*). Conversely, gliomas are immune-modulated despite being archetypically “cold”; even limited immune pressure shapes progression in the immune-privileged environment of the brain, as reflected by higher cytolytic activity in tumors relative to healthy tissue and evidence of immune evasion (*95, 122*). Taken together, these examples show how one-dimensional classifications flatten complex cancer biology into categories that may obscure how mutational and immune forces shape cancer incidence. By explicitly capturing this interplay, our framework provides an integrated basis for developing robust biomarkers.

While our analysis establishes a strong connection between immune decline and cancer risk, it also highlights key limitations and opportunities for future investigation. Longitudinal studies combining immunosequencing with clinical outcomes could help establish causal links. Repertoire-based analyses (*32, 123*–*126*) integrated with single-cell sequencing (*127*) may provide deeper insight into the cellular mechanisms of immunosenescence and how these processes differentially impact immune function. Signatures of immunoediting should be detectable in cancer genomes, yet broad pan-cancer analyses often find little signal once mutational covariates are controlled (*128, 129*). In contrast, studies of individual subtypes or single tumors report clear evidence of selection (*130*–*134*). This discrepancy, along with our results, suggests that immunoediting is context dependent, varying with host immune competence and cancer subtype. Our analysis identifies immune-modulated cancers as the settings where such signatures are most likely to be observed, implying that their detection depends on the underlying immune state of the host. Identifying robust measures of immune competence will be critical for understanding how immunity influences cancer risk, progression and therapeutic response. TCR diversity emerges as one such measure, associated both with cancer risk and with favorable outcomes and treatment responses (*135*–*142*), underscoring its translational potential.

We identify immune decline as a major contributor to cancer risk and show that cancer subtypes align along an immune-mutation axis that captures the joint influence of mutation and immunity. This framework formulates the concept of immunoediting as a mathematical model of carcinogenesis, linking immune competence to cancer risk through population-scale sequencing and epidemiological data. Understanding how immune variation shapes disease susceptibility, progression and treatment response will be key to reducing mortality. Our findings may guide advances in predictive modeling, immunoprevention, personalized medicine and next-generation immunotherapies—efforts urgently needed to confront the rising burden of cancer in an aging world.

## Materials & Methods

### Sequencing of Human Samples

Sequencing data from human samples used in this study were aggregated from a cohort of subjects who took the T-Detect Covid diagnostic test which received Emergency Use Authorization from the FDA. All human subjects provided informed consent and the study was approved by the WIRB Copernicus Group Institutional Review Board. Age and sex were recorded for all participants and the influence of these variables was explicitly analyzed and reported in the study.

A multi-plexed PCR method was used to sequence the CDR3 of TCR *β* chains of T cells, typically from 18*µ*g of genomic DNA (*59, 143*–*145*). Our results and conclusions are robust to small variations in the amount of input genomic DNA. Using the T-Detect Covid diagnostic test—which has high clinical performance (*145*)—we identify 9, 952 subjects in our sample with past or present SARS-CoV-2 infection, finding that past infections do not impact repertoire size or TCR diversity.

The median total number of productive TCRs sequenced (i.e., sequencing depth) for T-Detect Covid samples is 518,618 TCRs with 95% of subjects having a sequencing depth between 222,082 and 853,647. Self-identified males and females comprise 47.2% and 52.5% of subjects, respectively. The median age of the cohort is 50 years with 95% of subjects ranging between 20 and 74 years.

We estimate uncertainty in TCR diversity measurements from repeat measurements of the same subject. We identify 396 subjects with repeat samples on the basis of overlapping TCRs observed in their repertoires. Repeat samples from the same donor are identified by generating the Jaccard index of the 100 highest frequency clones in any two repertoires with a set threshold on the overlap. We further match these repeat samples on age and sex. The method is found to be 100% accurate in 446 samples, correctly categorizing these samples as belonging to 223 known unique donors. From repeat measurements of the same subject, we estimate a 1*σ* measurement uncertainty of 0.10 dex in the log of TCR diversity for a single sample (see Supplementary Material Figure S1). These random variations average out in our summary statistics.

TCRs are heterodimers and the same TCR*β* chain may randomly pair with different TCR*α*s on different T cells. Furthermore, TCR*β* repertoires significantly undersample the total number of unique clones in subjects, which is unknown, but is estimated to be *O*(10^7^) (*143*). However, we are primarily concerned with systematic changes and thus our analysis only requires *relative* accuracy in TCR diversity, which our data provide. Down-sampling repertoires to a fixed number of T cells yields qualitatively consistent results but introduces systematic bias due to prevalence of Cytomegalovirus infections (see Supplementary Material).

### Cancer Incidence Data

Cancer incidence data were retrieved via the SEER*Stat Software (*146*) from the Surveillance, Epidemiology and End Results (SEER) Program (www.seer.cancer.gov) SEER*Stat Database: Incidence - SEER Research Data, 22 Registries, Nov 2023 Sub (2000-2021) - Linked To County Attributes - Time Dependent (1990-2022) Income/Rurality, 1969-2022 Counties, National Cancer Institute, DCCPS, Surveillance Research Program, released April 2024, based on the November 2023 submission. We aggregate incidence data sorted by sex for all cancer types excluding sex-specific cancers (breast, cervix uteri, ovary, prostate, testis, uterus and vulva) and lung cancer due to known sex-linked behavioral differences (i.e., smoking). We restrict our analysis of cancer incidence to individuals between 20 and 80 years old.

In our subtype analysis, we restrict to cancers that satisfy the key model assumption that sex differences in incidence reflect differences in immune competence rather than sex bias due to anatomy, hormones, behavior or viral exposure. The set of 18 cancers subtypes that we include are: acute myeloid leukemia, chronic lymphocytic leukemia, chronic myeloid leukemia, colorectal adenocarcinoma, diffuse large B cell lymphoma, follicular lymphoma, gastric adenocarcinoma, kidney chromophobe carcinoma, kidney clear cell carcinoma, kidney papillary carcinoma, lung adenocarcinoma, lung squamous cell carcinoma, malignant gliomas (astrocytic tumors), melanoma of skin, multiple myeloma, pancreatic adenocarcinoma, lung small cell carcinoma and soft tissue sarcomas. Below we list exlcusion criteria.

Cancers excluded due to sex-specific anatomy include: prostate carcinoma, testicular germcell tumors, penile squamous cell carcinoma, ovarian carcinoma, endometrial (uterine) carcinoma (*147*), vaginal carcinoma and vulvar carcinoma (*148*). Female breast carcinoma (strong estrogen/progesterone signaling; large HR-positive fraction) (*149*) and papillary thyroid carcinoma (evidence for endocrine influence/ER signaling) (*150*) are excluded due to hormonally driven sex differences. Cancers excluded due to potentially sex-biased viral exposure-driven etiology include: HPV-associated oropharyngeal squamous cell carcinoma (*151*), HPV-associated anal squamous cell carcinoma (*152*), nasopharyngeal carcinoma (Epstein-Barr virus) (*153*), hepatocellular carcinoma (substantial HBV/HCV-attributable burden) (*154*) and Kaposi sarcoma (HHV-8/KSHV) (*155*). And those excluded due to behavioral/environmental exposure confounding include: laryngeal squamous cell carcinoma (tobacco-alcohol synergy) (*156*), oral-cavity squamous cell carcinoma (tobacco/alcohol) (*157*), hypopharyngeal/pharyngeal squamous cancers (tobacco/alcohol) (*158*), esophageal squamous cell carcinoma (tobacco/alcohol) (*159*), esophageal adenocarcinoma (GERD/obesity; marked male predominance) (*160*), bladder urothelial carcinoma (cigarette smoking and occupational aromatic amines) (*161*), sinonasal adenocarcinoma (wood/leather dust in predominantly male occupations) (*162*), malignant mesothelioma (asbestos) (*163*), basal cell carcinoma (UV radiation) (*164*) and cutaneous squamous cell carcinoma (UV radiation) (*165*). Finally, post-transplant lymphoproliferative disorders (PTLD) is excluded due to special-population immunosuppression (*166*).

### Literature data

We compare our fitted values of *γ* with independent estimates of driver mutation counts from Martincorena et al. (*92*), who used dN/dS analysis of TCGA sequences to infer the excess of nonsynonymous mutations under positive selection. Because no data table was provided, we visually extracted the relative ranking of cancers from their Figure 4A which showed the average number of driver mutations per tumor in 369 known cancer genes. We assessed concordance with our results using a Spearman rank correlation. Their extracted ordering (from high to low, using our naming convention) is: colorectal adenocarcinoma MSS, lung squamous cell carcinoma, lung adenocarcinoma, melanoma, glioblastoma, gastric adenocarcinoma, kidney clear cell carcinoma, soft tissue sarcoma and kidney papillary carcinoma.

### Intrinsic Scatter Calculation

We estimate the 1*σ* intrinsic scatter in TCR diversity, which appears to be log-normally distributed, as

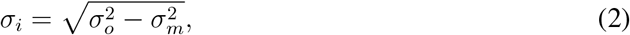

where *σ*_*i*_ is the intrinsic scatter, *σ*_*o*_ is the observed scatter containing the central 68% of samples and *σ*_*m*_ is the measurement uncertainty of 0.10 dex estimated from repeat samples. To estimate the intrinsic scatter at different quantiles of the distribution, we scale *σ*_*m*_ accordingly. For example to estimate the intrinsic scatter of the central 90% of samples, we calculate the half-confidence interval containing the central 90% of the data and subtract *σ*_*m*_ scaled by 1.645 in quadrature (for a normal distribution, 90% of samples are within *±*1.645*σ*) and multiply by 2.

### Age Offset Between Male and Female Repertoires

We derive the age offset between male and female repertoires (Figures 2C and 2D) and estimate statistical uncertainty by independently bootstrapping the male and female repertoires. For each bootstrapped realization, we generate the median repertoire size and TCR diversity (i.e., Figures 2A and 2B) as function of age and determine the age offset that minimizes the sum of squared residuals between the male and age-offset female curves. In order to match the age grid between male and age-offset female curves, we interpolate the curves for males and age-offset females between ages of 18 and 70 in 1 year increments. We limit the maximum interpolated age to 70 years to avoid significant extrapolation of the age-offset female curves. We report the mean and standard deviation of 10,001 realizations as the age offset and 1*σ* statistical uncertainty, respectively.

### Model Fitting

We fit cancer incidence as a function of age and TCR diversity using the Python routine curve fit implemented in the SciPy package (*167*). We input the standard errors of the cancer incidence data to weight the fit but derive parameter uncertainties via bootstrapping to account for the significantly larger uncertainties associated with our measurements of TCR diversity which can not be straightforwardly accounted for in standard non-linear least squares fitting. When fitting the cancer incidence we interpolate the median relation of *D*(*t*) for males and females separately (Figure 2B) using CubicSpline implemented in SciPy. Our results are insensitive to the polynomial degree of the spline. Interpolation of *D* is necessary because the age grid of *D* is not coincident with the cancer incidence data. Fitting a model to the log of cancer incidence yields consistent results.

We are unable to use standard error propagation because we have errors in both cancer incidence and TCR diversity data and the latter are treated as a dependent variable in our fit. To estimate the errors of our parameters, we bootstrap the repertoire data prior to generating the median *D*(*t*) for males and females (i.e., Figure 2B) via the binning procedure described in the text. Each bootstrapped realization is fit using the procedure described above. The dominant source of statistical uncertainty in our fit is from our measurement of *D* and our bootstrapping procedure effectively propagates these uncertainties to the model parameters.

We generate the 95% confidence interval of our fit shown in Figure 3A by propagating the 2*σ* uncertainties associated with *D* (see Table 1) through the best-fit model. Meaning, when generating the upper confidence interval, we evaluate our best-fit model adopting values of *D* that are 2*σ* larger, where *σ* is the error associated with each point in Figure 2B. We treat the lower bound similarly and interpolate *D* as before. Adopting 95% confidence from the bootstrapped fits yields similar intervals but fails to account for covariance.

For simplicity, we adopt the median TCR diversity as a function of age when fitting cancer incidence. One potential issue with using the median relation is that it does not account for the fact that in our model, individuals with lower TCR diversity have higher risk. We test whether adopting the median TCR diversity, rather than using TCR diversity of individual subjects, impacts our results. We calculate the expectation value for cancer incidence, ⟨*I*⟩, in each decade age bin as

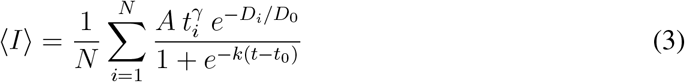

where *t*_*i*_ and *D*_*i*_ are age and TCR diversity for a given subject, respectively, and the sum is over all subjects in the age bin. *A, γ* and *D*_0_ are free parameters, as before. We calculate ⟨*I*⟩ for males and females independently and interpolate ⟨*I*⟩ across age bins to match the binning of the cancer incidence data. We shift ⟨*I*⟩ by 5 years to match age offset in Equation 1. This shift is meant to align the elimination phase we model with clinical detection. We use the same procedure described above to generate best-fit parameters and 95% confidence intervals yielding: 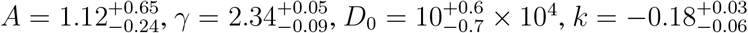 and 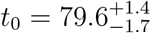. The use of individual measurements has a marginal effect on the parameters. Our results and conclusions are robust to use of median TCR diversity instead of individual measurements.

To assess parameter sensitivity, we varied the latency offset between 0 and 10 years (Figures S5 and S6). Increasing the latency offset systematically lowered the fitted values of both the normalization constant *A* and the mutational exponent *γ*, reflecting covariance among these parameters. The remaining parameters were consistent within statistical uncertainties. This behavior arises because larger latency values shift explanatory weight from the mutational component to the immune component of the model. Despite these covariances, the relative ordering of subtypes, their positions along the immune-mutation axis and the overall model goodness of fit remain stable, indicating that our conclusions are robust to reasonable variations in latency assumptions. The observed covariance underscores the difficulty of disentangling the accumulation of mutations from immune decline using incidence data alone, highlighting the need for longitudinal or mechanistic datasets.

### Partitioning Risk and Calculating TCR diversity percentiles

We partition risk into a component attributable to aging and one attributable to variation in TCR diversity at a given age. Risk due to aging is uniform within an age group and reflects contributions from both the accumulation of mutations and systematic immune decline. Mutation accumulation is modeled explicitly as a function of age, while risk associated with systematic age-related changes in TCR diversity is derived empirically from the median decline as a function of age (Figure 1B). In contrast, risk at a given age ranges among individuals due to differences in TCR diversity, which varies substantially (Figure 1B).

To partition the risk, we calculate cancer risk as a function of age and TCR diversity using our best-fit model and then relate TCR diversity at a given age to its percentile within the overall distribution of TCR diversity for that age group. For instance, while the median TCR diversity declines with age, it represents the 50th percentile of the TCR diversity distribution within any age group, meaning a given percentile retains the same relative position within the distribution despite the overall decline in TCR diversity with age. Thus, risk associated with both the accumulation of mutations and the general decline in TCR diversity (median curve in Figure 1B) is attributed to aging. In contrast, within any age group, variable risk results from differences in TCR diversity among individuals.

We derive the 1*σ* intrinsic scatter of log *D* as a function of age (Figure 1B) as described in the text. We then use this 1*σ* intrinsic scatter and assume a normal distribution to relate values of *D* to their corresponding percentiles which we then evaluate using our best fit model to generate a prediction of cancer incidence. Interpolation is necessary to generate the smooth map shown in Figure 3B and introduces minimal errors that do not impact our conclusions. To estimate relative risk, we normalize all predictions to a 40 year-old subject with median TCR diversity.

## Data Availability

Data table with T cell repertoire metrics available at https://doi.org/10.5281/zenodo.13993996.

## Acknowledgments

We thank Mary Carrington, Margaret Geller, Matthew Noakes, Ash Chapfuwa, Miah Wander and Ko-Han Lee for reading the manuscript and providing constructive comments, Beryl Crossley for developing the method to identify repeat samples from the same donor and Rebecca Elyanow for its implementation in our software pipeline.

## Competing Interests

HJ Zahid, J Greissl have employment and equity ownership with Microsoft. MG Noceda, HS Robins have employment and equity ownership with Adaptive Biotechnologies. The authors declare no other competing interests.

During the preparation of this work HJZ used ChatGPT in order to assist in editing the manuscript for clarity and readability. After using this tool/service, HJZ reviewed and edited the content as needed and takes full responsibility for the content of the publication.

## Footnotes

1 We use cancer incidence and cancer risk interchangeably throughout, as the distinction between the two is minimal in the context of first-time cancer diagnoses and does not impact our analysis (*3*).

2 https://www.fda.gov/media/146481/download

3 A difference of *x* dex indicates a change by a factor of 10^*x*^.

## References

(1) Kanasi, E., Ayilavarapu, S., and Jones, J. (2016). The aging population: demographics and the biology of aging. Periodontology 2000 72, 13–18.

(2) J, F. et al. Global Cancer Observatory: Cancer Tomorrow (version 1.1) ed. by for Research on Cancer, I. A., https://gco.iarc.who.int/tomorrow (accessed Sept. 10, 2024).

(3) Sasieni, P. et al. (2011). What is the lifetime risk of developing cancer?: the effect of adjusting for multiple primaries. British journal of cancer 105, 460–465.

(4) Ehrlich, P., Ueber den jetzigen Stand der Karzinomforschung, 1908.

(5) Burnet, M., Immunological surveillance; Elsevier: 2014.

(6) Dunn, G. P. et al. (2002). Cancer immunoediting: from immunosurveillance to tumor escape. Nature immunology 3, 991–998.

(7) Schreiber, R. D., Old, L. J., and Smyth, M. J. (2011). Cancer immunoediting: integrating immunity’s roles in cancer suppression and promotion. Science 331, 1565–1570.

(8) Vesely, M. D. et al. (2011). Natural innate and adaptive immunity to cancer. Annual review of immunology 29, 235–271.

(9) Schumacher, T. N., Scheper, W., and Kvistborg, P. (2019). Cancer neoantigens. Annual review of immunology 37, 173–200.

(10) Hedrick, S. M. et al. (1984). Isolation of cDNA clones encoding T cell-specific membrane-associated proteins. Nature 308, 149–153.

(11) Yanagi, Y. et al. (1984). A human T cell-specific cDNA clone encodes a protein having extensive homology to immunoglobulin chains. Nature 308, 145–149.

(12) Davis, M. M. et al. (1998). Ligand recognition by (alpha)(beta) T cell receptors. Annual review of immunology 16, 523.

(13) Nikolich-Žugich, J., Slifka, M. K., and Messaoudi, I. (2004). The many important facets of T-cell repertoire diversity. Nature Reviews Immunology 4, 123–132.

(14) Foster, A. D., Sivarapatna, A., and Gress, R. E. (2011). The aging immune system and its relationship with cancer. Aging health 7, 707–718.

(15) Martin, M. P., and Carrington, M. (2013). Immunogenetics of HIV disease. Immunological reviews 254, 245–264.

(16) Corthay, A. (2014). Does the immune system naturally protect against cancer? Frontiers in immunology 5, 197.

(17) Montgomery, R. A. et al. (2018). HLA in transplantation. Nature reviews nephrology 14, 558–570.

(18) Kovacs, A. A. et al. (2020). Association of HLA Genotype With T-Cell Activation in Human Immunodeficiency Virus (HIV) and HIV/Hepatitis C Virus–Coinfected Women. The Journal of infectious diseases 221, 1156–1166.

(19) Francis, J. M. et al. (2021). Allelic variation in class I HLA determines CD8+ T cell repertoire shape and cross-reactive memory responses to SARS-CoV-2. Science immunology 7, eabk3070.

(20) Granadier, D. et al. In Seminars in immunopathology, 2021; Vol. 43, pp 119–134.

(21) Olafsdottir, T. A. et al. (2022). HLA alleles, disease severity, and age associate with T-cell responses following infection with SARS-CoV-2. Communications Biology 5, 914.

(22) Hozumi, N., and Tonegawa, S. (1976). Evidence for somatic rearrangement of immunoglobulin genes coding for variable and constant regions. Proceedings of the National Academy of Sciences 73, 3628–3632.

(23) Aspinall, R., and Andrew, D. (2000). Thymic involution in aging. Journal of clinical immunology 20, 250–256.

(24) Walford, R. L. (1964). The immunologic theory of aging. The Gerontologist 4, 195–197.

(25) Aw, D., Silva, A. B., and Palmer, D. B. (2007). Immunosenescence: emerging challenges for an ageing population. Immunology 120, 435–446.

(26) Gruver, A., Hudson, L., and Sempowski, G. (2007). Immunosenescence of ageing. The Journal of Pathology: A Journal of the Pathological Society of Great Britain and Ireland 211, 144–156.

(27) Pawelec, G. (2018). Age and immunity: what is “immunosenescence”? Experimental gerontology 105, 4–9.

(28) Mittelbrunn, M., and Kroemer, G. (2021). Hallmarks of T cell aging. Nature immunology 22, 687–698.

(29) Liu, Z. et al. (2023). Immunosenescence: molecular mechanisms and diseases. Signal transduction and targeted therapy 8, 200.

(30) Naylor, K. et al. (2005). The influence of age on T cell generation and TCR diversity. The Journal of Immunology 174, 7446–7452.

(31) Goronzy, J. J., Lee, W.-W., and Weyand, C. M. (2007). Aging and T-cell diversity. Experimental gerontology 42, 400–406.

(32) Qi, Q. et al. (2014). Diversity and clonal selection in the human T-cell repertoire. Proceedings of the National Academy of Sciences 111, 13139–13144.

(33) Britanova, O. V. et al. (2014). Age-related decrease in TCR repertoire diversity measured with deep and normalized sequence profiling. The Journal of Immunology 192, 2689–2698.

(34) Goronzy, J. J., and Weyand, C. M. (2017). Successful and maladaptive T cell aging. Immunity 46, 364–378.

(35) Fulop, T. et al. (2010). Potential role of immunosenescence in cancer development. Annals of the new York Academy of Sciences 1197, 158–165.

(36) Goronzy, J. J., and Weyand, C. M. (2013). Understanding immunosenescence to improve responses to vaccines. Nature immunology 14, 428–436.

(37) Gil, A. et al. (2015). Narrowing of human influenza A virus-specific T cell receptor α and β repertoires with increasing age. Journal of virology 89, 4102–4116.

(38) Pinti, M. et al. (2016). Aging of the immune system: focus on inflammation and vaccination. European journal of immunology 46, 2286–2301.

(39) Del Giudice, G. et al. (2017). Fighting against a protean enemy: immunosenescence, vaccines, and healthy aging. npj Aging and Mechanisms of Disease 4, 1.

(40) Liu, W. et al. (2020). Analysis of factors associated with disease outcomes in hospitalized patients with 2019 novel coronavirus disease. Chinese medical journal 133, 1032–1038.

(41) Zhou, F. et al. (2020). Clinical course and risk factors for mortality of adult inpatients with COVID-19 in Wuhan, China: a retrospective cohort study. The lancet 395, 1054–1062.

(42) Klein, S. L., and Flanagan, K. L. (2016). Sex differences in immune responses. Nature Reviews Immunology 16, 626–638.

(43) Van Lunzen, J., and Altfeld, M. (2014). Sex differences in infectious diseases–common but neglected. The Journal of infectious diseases 209, S79–S80.

(44) vom Steeg, L. G., and Klein, S. L. (2016). SeXX matters in infectious disease pathogenesis. PLoS pathogens 12, e1005374.

(45) Klein, S. L., Jedlicka, A., and Pekosz, A. (2010). The Xs and Y of immune responses to viral vaccines. The Lancet infectious diseases 10, 338–349.

(46) Cook, M. B. et al. (2009). Sex disparities in cancer incidence by period and age. Cancer Epidemiology Biomarkers & Prevention 18, 1174–1182.

(47) Dong, M. et al. (2020). Sex differences in cancer incidence and survival: a pan-cancer analysis. Cancer Epidemiology, Biomarkers & Prevention 29, 1389–1397.

(48) Yan, J. et al. (2010). The effect of ageing on human lymphocyte subsets: comparison of males and females. Immunity & Ageing 7, 1–10.

(49) Kverneland, A. H. et al. (2016). Age and gender leucocytes variances and references values generated using the standardized ONE-Study protocol. Cytometry part A 89, 543–564.

(50) Pido-Lopez, J., Imami, N., and Aspinall, R. (2001). Both age and gender affect thymic output: more recent thymic migrants in females than males as they age. Clinical & Experimental Immunology 125, 409–413.

(51) Stankiewicz, L. N. et al. (2025). Sex-biased human thymic architecture guides T cell development through spatially defined niches. Developmental Cell 60, 152–169.

(52) Hahn, P. A. et al. (2025). Exogenous estrogen enhances T cell activation in male primates. Cell Reports 44.

(53) Armitage, P., and Doll, R. (2004). The age distribution of cancer and a multi-stage theory of carcinogenesis. British journal of cancer 91, 1983–1989.

(54) Vineis, P., Schatzkin, A., and Potter, J. D. (2010). Models of carcinogenesis: an overview. Carcinogenesis 31, 1703–1709.

(55) Palmer, S. et al. (2018). Thymic involution and rising disease incidence with age. Proceedings of the National Academy of Sciences 115, 1883–1888.

(56) Liu, X., and Wu, J. (2018). History, applications, and challenges of immune repertoire research. Cell biology and toxicology 34, 441–457.

(57) Bradley, P., and Thomas, P. G. (2019). Using T cell receptor repertoires to understand the principles of adaptive immune recognition. Annual review of immunology 37, 547–570.

(58) Cowell, L. G. (2020). The Diagnostic, Prognostic, and Therapeutic Potential of Adaptive Immune Receptor Repertoire Profiling in CancerAdaptive Immune Receptor Repertoire Profiling in Cancer. Cancer research 80, 643–654.

(59) Carlson, C. S. et al. (2013). Using synthetic templates to design an unbiased multiplex PCR assay. Nature communications 4, 1–9.

(60) Robins, H. (2013). Immunosequencing: applications of immune repertoire deep sequencing. Current opinion in immunology 25, 646–652.

(61) Sender, R. et al. (2023). The total mass, number, and distribution of immune cells in the human body. Proceedings of the National Academy of Sciences 120, e2308511120.

(62) Lang, P. O., Govind, S., and Aspinall, R. (2013). Reversing T cell immunosenescence: why, who, and how. Age 35, 609–620.

(63) Duggal, N. A. (2018). Reversing the immune ageing clock: lifestyle modifications and pharmacological interventions. Biogerontology 19, 481–496.

(64) Aspinall, R., and Lang, P. O. (2018). Interventions to restore appropriate immune function in the elderly. Immunity & Ageing 15, 1–8.

(65) Lanfermeijer, J., Borghans, J. A., and van Baarle, D. (2020). How age and infection history shape the antigen-specific CD8+ T-cell repertoire: Implications for vaccination strategies in older adults. Aging Cell 19, e13262.

(66) Zahid, H. J. et al. (2025). A Fundamental Relationship between TCR Diversity, Repertoire Size and Systemic Clonal Expansion: Insights from 30,000 TCRβ Repertoires. bioRxiv, 2025–05.

(67) Poisner, H. et al. (2024). Genetic determinants and phenotypic consequences of blood T-cell proportions in 207,000 diverse individuals. Nature Communications 15, 6732.

(68) Bentham, R. et al. (2025). ImmuneLENS characterizes systemic immune dysregulation in aging and cancer. Nature Genetics, 1–12.

(69) Manuel, M. et al. (2012). Lymphopenia combined with low TCR diversity (divpenia) predicts poor overall survival in metastatic breast cancer patients. Oncoimmunology 1, 432–440.

(70) Charles, J. et al. (2020). T-cell receptor diversity as a prognostic biomarker in melanoma patients. Pigment cell & melanoma research 33, 612–624.

(71) Kim, D. W. et al. Increased T-cell receptor repertoire diversity to predict better overall survival in gastrointestinal malignancies. 2021.

(72) Porciello, N. et al. (2022). T-cell repertoire diversity: friend or foe for protective antitumor response? Journal of Experimental & Clinical Cancer Research 41, 356.

(73) Wiegel, F. W., and Perelson, A. S. (2004). Some scaling principles for the immune system. Immunology and cell biology 82, 127–131.

(74) Zarnitsyna, V. I. et al. (2013). Estimating the diversity, completeness, and cross-reactivity of the T cell repertoire. Frontiers in immunology 4, 485.

(75) Harding, C., Pompei, F., and Wilson, R. (2012). Peak and decline in cancer incidence, mortality, and prevalence at old ages. Cancer 118, 1371–1386.

(76) Pavlidis, N., Stanta, G., and Audisio, R. A. (2012). Cancer prevalence and mortality in centenarians: a systematic review. Critical reviews in oncology/hematology 83, 145–152.

(77) Yachida, S. et al. (2010). Distant metastasis occurs late during the genetic evolution of pancreatic cancer. Nature 467, 1114–1117.

(78) Brenner, H. et al. (2011). Sojourn time of preclinical colorectal cancer by sex and age: estimates from the German national screening colonoscopy database. American Journal of Epidemiology 174, 1140–1146.

(79) Nadler, D. L., and Zurbenko, I. G. (2014). Estimating Cancer Latency Times Using a Weibull Model. Advances in Epidemiology 2014, 746769.

(80) Gerstung, M. et al. (2020). The evolutionary history of 2,658 cancers. Nature 578, 122–128.

(81) Grulich, A. E. et al. (2007). Incidence of cancers in people with HIV/AIDS compared with immunosuppressed transplant recipients: a meta-analysis. The Lancet 370, 59–67.

(82) Ilham, S. et al. (2023). Cancer incidence in immunocompromised patients: a single-center cohort study. BMC cancer 23, 33.

(83) Kooshesh, K. A. et al. (2023). Health consequences of thymus removal in adults. New England Journal of Medicine 389, 406–417.

(84) Khuder, S. A. (2001). Effect of cigarette smoking on major histological types of lung cancer: a meta-analysis. Lung cancer 31, 139–148.

(85) Thun, M. J. et al. (2013). 50-year trends in smoking-related mortality in the United States. New England Journal of Medicine 368, 351–364.

(86) Islami, F., Torre, L. A., and Jemal, A. (2015). Global trends of lung cancer mortality and smoking prevalence. Translational lung cancer research 4, 327.

(87) Vajdic, C. M. et al. (2007). Cancer incidence before and after kidney transplantation. The Lancet 370, 1871–1878.

(88) Engels, E. A. et al. (2011). Spectrum of cancer risk among US solid organ transplant recipients. JAMA 306, 1891–1901.

(89) Hernández-Ramírez, R. U. et al. (2017). Cancer risk in HIV-infected people in the USA from 1996 to 2012: a population-based, registry-linkage study. The Lancet HIV 4, e495–e504.

(90) Mayor, P. C. et al. (2018). Cancer in primary immunodeficiency diseases: Cancer incidence in the US Immune Deficiency Network registry. Journal of Allergy and Clinical Immunology 141, 1028–1035.

(91) Jackson, S. S. et al. (2024). Sex differences in cancer incidence among solid organ transplant recipients. JNCI: Journal of the National Cancer Institute 116, 401–407.

(92) Martincorena, I. et al. (2017). Universal patterns of selection in cancer and somatic tissues. Cell 171, 1029–1041.

(93) Tomasetti, C., and Vogelstein, B. (2015). Variation in cancer risk among tissues can be explained by the number of stem cell divisions. Science 347, 78–81.

(94) Tomasetti, C., Li, L., and Vogelstein, B. (2017). Stem cell divisions, somatic mutations, cancer etiology, and cancer prevention. Science 355, 1330–1334.

(95) Rooney, M. S. et al. (2015). Molecular and genetic properties of tumors associated with local immune cytolytic activity. Cell 160, 48–61.

(96) Chen, D. S., and Mellman, I. (2017). Elements of cancer immunity and the cancer– immune set point. Nature 541, 321–330.

(97) Landau, D. A. et al. (2013). Evolution and impact of subclonal mutations in chronic lymphocytic leukemia. Cell 152, 714–726.

(98) Okosun, J. et al. (2014). Integrated genomic analysis identifies recurrent mutations and evolution patterns driving the initiation and progression of follicular lymphoma. Nature Genetics 46, 176–181.

(99) Bolli, N. et al. (2014). Heterogeneity of genomic evolution and mutational profiles in multiple myeloma. Nature Communications 5, 2997.

(100) Papaemmanuil, E., Gerstung, M., Bullinger, L., et al. (2016). Genomic classification and prognosis in acute myeloid leukemia. New England Journal of Medicine 374, 2209–2221.

(101) Georgiou, K. et al. (2016). Genomic basis of PD-L1 overexpression in diffuse large B-cell lymphomas. Blood 127, 3026–3034.

(102) Kataoka, K. et al. (2019). Aberrant PD-L1 expression through 3’-UTR disruption in multiple cancers. Cancer Discovery 9, 462–477.

(103) Dufva, O. et al. (2020). Immunogenomic Landscape of Hematological Malignancies. Cancer Cell 38, 380–399.e13.

(104) Ai, Z. et al. (2024). VISTA blockade synergizes with tyrosine kinase inhibitors in chronic myeloid leukemia by enhancing T cell activity. Biomolecules & Therapeutics 32, e2024017.

(105) Davis, C. F. et al. (2014). The Somatic Genomic Landscape of Chromophobe Renal Cell Carcinoma. Cancer Cell 26, 319–330.

(106) Mitchell, T. J. et al. (2018). Genomics and clinical correlates of renal cell carcinoma. World Journal of Urology 36, 1899–1911.

(107) Braun, D. A. et al. (2020). Interplay of somatic alterations and immune infiltration modulates response to PD-1 blockade in advanced clear cell renal cell carcinoma. Nature Medicine 26, 909–918.

(108) George, J. et al. (2015). Comprehensive genomic profiles of small cell lung cancer. Nature 524, 47–53.

(109) Alexandrov, L. B. et al. (2016). Mutational signatures associated with tobacco smoking in human cancer. Science 354, 618–622.

(110) de Martel, C. et al. (2020). Global burden of cancer attributable to infections in 2018: a worldwide incidence analysis. The Lancet global health 8, e180–e190.

(111) Bentham, R. et al. (2021). Using DNA sequencing data to quantify T cell fraction and therapy response. Nature 597, 555–560.

(112) Bailey, P. et al. (2016). Genomic analyses identify molecular subtypes of pancreatic cancer. Nature 531, 47–52.

(113) Hodis, E. et al. (2012). A landscape of driver mutations in melanoma. Cell 150, 251–263.

(114) Snyder, A. et al. (2014). Genetic basis for clinical response to CTLA-4 blockade in melanoma. New England Journal of Medicine 371, 2189–2199.

(115) Rizvi, N. A. et al. (2015). Mutational landscape determines sensitivity to PD-1 blockade in non–small cell lung cancer. Science 348, 124–128.

(116) Chan, T. A. et al. (2019). Development of tumor mutation burden as an immunotherapy biomarker: utility for the oncology clinic. Cancer Cell 36, 591–595.

(117) Marcus, L. et al. (2021). FDA Approval Summary: Pembrolizumab for the Treatment of Tumor Mutational Burden–High Solid Tumors. Clinical Cancer Research 27, 4685–4689.

(118) McGrail, D. J. et al. (2021). High tumor mutation burden fails to predict immune checkpoint blockade response across all cancer types. Annals of Oncology 32, 661–672.

(119) Zheng, M. et al. (2022). Tumor mutational burden as a predictive biomarker for immunotherapy in solid tumors: a systematic review and meta-analysis. Frontiers in Oncology 12, 820418.

(120) Network, C. G. A. et al. (2012). Comprehensive molecular characterization of human colon and rectal cancer. Nature 487, 330.

(121) Angelova, M. et al. (2015). Characterization of the immunophenotypes and antigenomes of colorectal cancers reveals distinct tumor escape mechanisms and novel targets for immunotherapy. Genome Biology 16, 64.

(122) Sampson, J. H. et al. (2020). Brain immunology and immunotherapy in brain tumours. Nature Reviews Cancer 20, 12–25.

(123) Emerson, R. O. et al. (2017). Immunosequencing identifies signatures of cytomegalovirus exposure history and HLA-mediated effects on the T cell repertoire. Nature genetics 49, 659–665.

(124) DeWitt III, W. S. et al. (2018). Human T cell receptor occurrence patterns encode immune history, genetic background, and receptor specificity. Elife 7, e38358.

(125) Greissl, J. et al. (2021). Immunosequencing of the T-cell receptor repertoire reveals signatures specific for diagnosis and characterization of early Lyme disease. medRxiv.

(126) Zahid, H. J. et al. (2025). Large-scale statistical mapping of T-cell receptor β sequences to human leukocyte antigens. Frontiers in Immunology Volume 16 - 2025, DOI: 10.3389/fimmu.2025.1603730.

(127) Mogilenko, D. A., Shchukina, I., and Artyomov, M. N. (2022). Immune ageing at single-cell resolution. Nature Reviews Immunology 22, 484–498.

(128) Van den Eynden, J. et al. (2019). Lack of detectable neoantigen depletion signals in the untreated cancer genome. Nature Genetics 51, 1741–1748.

(129) Claeys, A. et al. (2021). Low immunogenicity of common cancer hot spot mutations resulting in false immunogenic selection signals. PLoS Genetics 17, e1009368.

(130) Marty, R. et al. (2017). MHC-I genotype restricts the oncogenic mutational landscape. Cell 171, 1272–1283.e15.

(131) Marty, R. et al. (2018). Evolutionary pressure against MHC class II binding cancer mutations. Cell 175, 416–428.e13.

(132) Zhang, A. W. et al. (2018). Interfaces of malignant and immunologic clonal dynamics in ovarian cancer. Cell 173, 1755–1769.e22.

(133) Rosenthal, R. et al. (2019). Neoantigen-directed immune escape in lung cancer evolution. Nature 567, 479–485.

(134) Łuksza, M. et al. (2022). Neoantigen quality predicts immunoediting in survivors of pancreatic cancer. Nature 606, 389–395.

(135) Postow, M. A. et al. (2015). Peripheral T cell receptor diversity is associated with clinical outcomes following ipilimumab treatment in metastatic melanoma. Journal for immunotherapy of cancer 3, 1–5.

(136) Kansy, B. A. et al. (2018). T cell receptor richness in peripheral blood increases after cetuximab therapy and correlates with therapeutic response. Oncoimmunology 7, e1494112.

(137) Guo, L. et al. (2020). Characteristics, dynamic changes, and prognostic significance of TCR repertoire profiling in patients with renal cell carcinoma. The Journal of pathology 251, 26–37.

(138) Han, J. et al. (2020). TCR repertoire diversity of peripheral PD-1+ CD8+ T cells predicts clinical outcomes after immunotherapy in patients with non–small cell lung cancer. Cancer immunology research 8, 146–154.

(139) Milano, F. et al. (2020). Impact of T cell repertoire diversity on mortality following cord blood transplantation. Frontiers in Oncology 10, 583349.

(140) Cardinale, A. et al. (2021). Thymic function and T-cell receptor repertoire diversity: implications for patient response to checkpoint blockade immunotherapy. Frontiers in immunology 12, 752042.

(141) Altan, M. et al. (2024). High peripheral T cell diversity is associated with lower risk of toxicity and superior response to dual immune checkpoint inhibitor therapy in patients with metastatic NSCLC. Journal for immunotherapy of cancer 12, e008950.

(142) Kjær, A. et al. (2025). Low T cell diversity associates with poor outcome in bladder cancer: A comprehensive longitudinal analysis of the T cell receptor repertoire. Cell Reports Medicine 6.

(143) Robins, H. S. et al. (2009). Comprehensive assessment of T-cell receptor β-chain diversity in αβ T cells. Blood, The Journal of the American Society of Hematology 114, 4099–4107.

(144) Robins, H. et al. (2012). Ultra-sensitive detection of rare T cell clones. Journal of immunological methods 375, 14–19.

(145) Dalai, S. C. et al. (2022). Clinical validation of a novel T-cell receptor sequencing assay for identification of recent or prior severe acute respiratory syndrome coronavirus 2 infection. Clinical Infectious Diseases 75, 2079–2087.

(146) National Cancer Institute SEER*Stat, version 8.4.3, 2024.

(147) Morice, P. et al. (2016). Endometrial cancer. The Lancet 387, 1094–1108.

(148) Preti, M. et al. (2005). Vulvar intraepithelial neoplasia. Best Practice & Research Clinical Obstetrics & Gynaecology 19, 591–602.

(149) Harbeck, N., and Gnant, M. (2017). Breast cancer. The Lancet 389, 1134–1150.

(150) Rahbari, R., Zhang, L., and Kebebew, E. (2010). Thyroid cancer gender disparity. Future Oncology 6, 1771–1779.

(151) Chaturvedi, A. K. et al. (2011). Human papillomavirus and rising oropharyngeal cancer incidence in the United States. Journal of Clinical Oncology 29, 4294–4301.

(152) de Martel, C. et al. (2017). Worldwide burden of cancer attributable to HPV by site, country and HPV type. International Journal of Cancer 141, 664–670.

(153) Chen, Y.-P. et al. (2019). Nasopharyngeal carcinoma. The Lancet 394, 64–80.

(154) Llovet, J. M. et al. (2021). Hepatocellular carcinoma. Nature Reviews Disease Primers 7, 6.

(155) Cesarman, E. et al. (2019). Kaposi sarcoma. Nature Reviews Disease Primers 5, 9.

(156) Hashibe, M. et al. (2009). Interaction between tobacco and alcohol use and the risk of head and neck cancer: pooled analysis in the International Head and Neck Cancer Epidemiology Consortium. Cancer Epidemiology, Biomarkers & Prevention 18, 541–550.

(157) Blot, W. J. et al. (1988). Smoking and drinking in relation to oral and pharyngeal cancer. Cancer Research 48, 3282–3287.

(158) Hashibe, M. et al. (2007). Alcohol drinking in never users of tobacco, cigarette smoking in never drinkers, and the risk of head and neck cancer: pooled analysis in the International Head and Neck Cancer Epidemiology Consortium. Journal of the National Cancer Institute 99, 777–789.

(159) Abnet, C. C., Arnold, M., and Wei, W.-Q. (2018). Epidemiology of esophageal squamous cell carcinoma. Gastroenterology 154, 360–373.

(160) Thrift, A. P., and Whiteman, D. C. (2012). The incidence of esophageal adenocarcinoma continues to rise: analysis of period and birth cohort effects on recent trends. Annals of Oncology 23, 3155–3162.

(161) Cumberbatch, M. G. et al. (2016). The role of tobacco smoke in bladder and kidney carcinogenesis: a comparison of exposures and meta-analysis of incidence and mortality risks. European Urology 70, 458–466.

(162) Leclerc, A. et al. (1994). Sino-nasal cancer and wood dust exposure: Results from a case-control study. American Journal of Epidemiology 140, 340–349.

(163) Carbone, M. et al. (2019). Mesothelioma: scientific clues for prevention, diagnosis, and therapy. CA: A Cancer Journal for Clinicians 69, 402–429.

(164) Lomas, A., Leonardi-Bee, J., and Bath-Hextall, F. (2012). A systematic review of worldwide incidence of nonmelanoma skin cancer. British Journal of Dermatology 166, 1069–1080.

(165) Karia, P. S., Han, J., and Schmults, C. D. (2013). Cutaneous squamous cell carcinoma: estimated incidence of disease, nodal metastasis, and deaths from disease in the United States, 2012. Journal of the American Academy of Dermatology 68, 957–966.

(166) Dharnidharka, V. R. et al. (2016). Post-transplant lymphoproliferative disorders. Nature Reviews Disease Primers 2, 15088.

(167) Virtanen, P. et al. (2020). SciPy 1.0: Fundamental Algorithms for Scientific Computing in Python. Nature Methods 17, 261–272.

