## Supplemental Material for "The Immune-Mutation Axis of Cancer Incidence: Insights from 30,000 TCR*β* Repertoires"

### Supplementary Material

#### S1. Statistical uncertainty in TCR diversity from repeat measurements

Figure S1 shows the distribution of differences in TCR diversity derived from 396 repeat samples from the same subjects. We estimate a statistical uncertainty of $\sim 0.10$ dex associated with the measurement of TCR diversity from a single repertoire. This estimate accounts for both random variations in the measurement and some short-term temporal fluctuations since repeat samples are not always from contemporaneous blood draws. Thus, it may slightly overestimate uncertainty associated solely with sequencing.

#### S2. T cell diversity and its dependence on sex in an independent cohort

We test results presented in the main text using an independent cohort. Whole blood samples from DLS (Discovery Life Sciences, Huntsville, AL) were collected under Protocol DLS13 for collection of clinical samples. Samples were sequenced using the same assay as the T-Detect Covid samples but subjects are completely distinct in the two samples. Figure S2 reproduces Figure 2 in the main text using 1734 independent samples. In these data, the female curves translate into the male curves shifting $16.0 \pm 2.5$ years which is statistically consistent ($1.7\sigma$) with $11.4 \pm 0.9$ years in the main text; differences in diversity between males and females in this cohort appear at younger ages as compared to the T-Detect COVID cohort analyzed in the main text. However, the primary conclusion of Figure 2—that females have higher TCR diversity such that their repertoires are consistent with males that are younger—and results of subsequent model analysis are unchanged because our model is primarily constrained by relative differences in diversity at older ages when sex-linked disparities in cancer incidence become pronounced and thus are insensitive to the younger age at which diversity begins to deviate between males and females in the DLS cohort.

#### S3. CMV positivity, T cell diversity and down-sampled repertoires

An alternative to measuring TCR diversity directly from sequenced repertoires is to sub-sample repertoires to some fiducial sequencing depth to account for potential systematic differences in the sequencing procedure, e.g., (*1*). This approach provides a measure of T cell diversity per fixed number of sequenced cells and is useful if systematic variations in sequencing depth are large. For instance, if repertoire data is from different cohorts which are not homogeneously sequenced. However, a challenge of this procedure is that it is difficult to interpret the normalized TCR diversity due to systematic variations in repertoire size between subjects. For example, both age and CMV status, among other possible variables, impact repertoire size.

CMV is a chronic herpes-virus with high global seroprevalence and significant geographical variation (*2*). CMV can not be cleared from the body but is generally suppressed by the immune system of healthy individuals (*3*). Because of its chronic and persistent nature, CMV has a substantial impact on the T cell repertoire with significant clonal expansion of CMV specific T cells, particularly CD8$^+$ T cells which can have oligiclonal populations that comprise a quarter of the T cell population (*4–6*). CMV has been linked to immunosenescence which may contribute to poorer health outcomes in the elderly though the impact is not well understood (*7*, *8*). Conversely, diversity of the T cell repertoire rather than size may be more important for properly functioning immune system (*9*) and studies suggest that CMV positive subjects maintain T cell diversity by increasing the size of their repertoire to accommodate expansion of CMV specific T cells (*10–12*).

We apply a sensitive and specific T cell based diagnostic for CMV to all repertoires we analyze using a method previously described in (*13–15*), which selects disease associated TCRs based on labeled case/control data. We use 2181 labeled samples from the DLS dataset described above with serological CMV labels to build a CMV classifier with an area under the receiver operating characteristic curve (AUROC) of 0.96 on the same holdout used in (*13*). This allows us to label the CMV status of all repertoires we analyze in this study and explore the impact of CMV postivity on repertoire size and diversity and subsequent systematics associated with down-sampling.

Figure S3 shows TCR diversity measured from down-sampled T-Detect Covid repertoires which all have sequencing depth greater than 50,000 T cells. Here we have randomly selected 50,000 TCRs from each repertoire and calculated the total number of unique TCRs, $D_{50}$. The results are qualitatively consistent with those based on measurements from the full repertoire as shown in the manuscript. Using only CMV negative subjects, we show that down-sampled TCR diversity declines with age (Figure S3A) and females appear to have greater diversity as compared to males (Figure S3B). Figure S3C shows a significant difference in down-sampled TCR diversity as a function of CMV status.

CMV has an enormous impact on TCR diversity because systematic changes in repertoire size associated with CMV positivity. Figure S4A shows that the total number of T cells sequenced is substantially larger in CMV positive subjects as compared to CMV negative subjects. This difference in size increases as a function of age. On the contrary, CMV has a small, albeit systematic, impact on the diversity of the T cell repertoire (Figure S4B). We estimate the relative increase in repertoire size in Figure S4C. CMV positive subjects at age 20 have a repertoire that is $\sim 5\%$ larger on average; the difference increases to $\sim 25\%$ by age 80. CMV persistently challenges the immune system and we attribute the increase in repertoire size to clonal expansion of CMV specific T cells. CMV prevalence increases as a function of age and thus the time since initial infection varies for subjects at any given time point in Figure S4C. The increase as a function of age thus results from a general increase in the clonality of CMV specific T cells but does not quantify the expected change in the repertoire size of any single subject as a function of time.

CMV has a significant impact on the size of T cell repertoire but the relatively small impact on TCR diversity is a consequence of the fact that the T cell repertoire grows larger to accommodate CMV specific T cells. As a consequence, TCR diversity measured from down-sampled repertoires will grossly underestimate the true diversity of CMV positive subjects because repertoires down-sampled to a fixed number of cells can not account for the increase in the total number of T cells in the body. Given the high prevelance of CMV in the population, this systematic effect introduces significant bias which must be accounted for if down-sampled repertoires are used to measure TCR diversity. In our analysis, measuring TCR diversity directly from the sequenced repertoire accounts for the systematically larger repertoire size resulting from CMV.

#### S4. Parameter Sensitivity Analysis

Figures S5 and S6 illustrate covariance between the latency parameter ($t_{\mathrm{off}}$) and the fitted parameters $D_0$ and $\gamma$, respectively, obtained from sensitivity analyses.

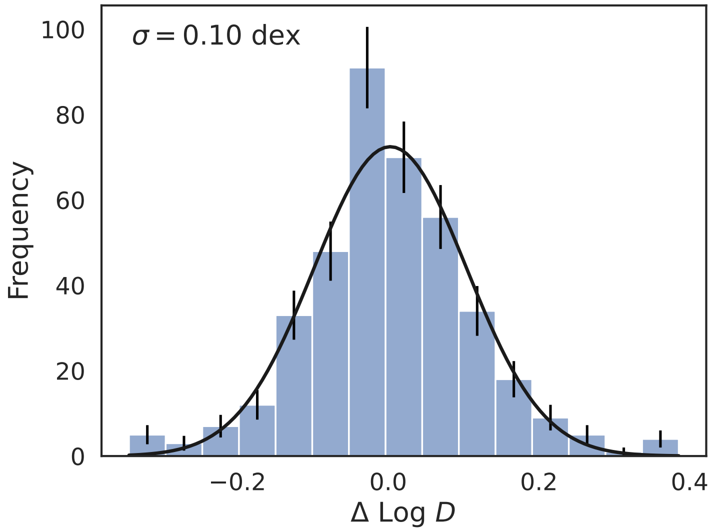


Figure S1: **Difference in TCR diversity from repeat observations of the same subject.** A histogram showing the distribution of differences in TCR diversity measured from 396 subjects with repeat samples. We fit the distribution with a gaussian using curve_fit in the SciPy package and estimate a standard deviation, $\sigma = 0.10$ dex. The data are close to log-normally distributed though there is an excess at small values near zero.

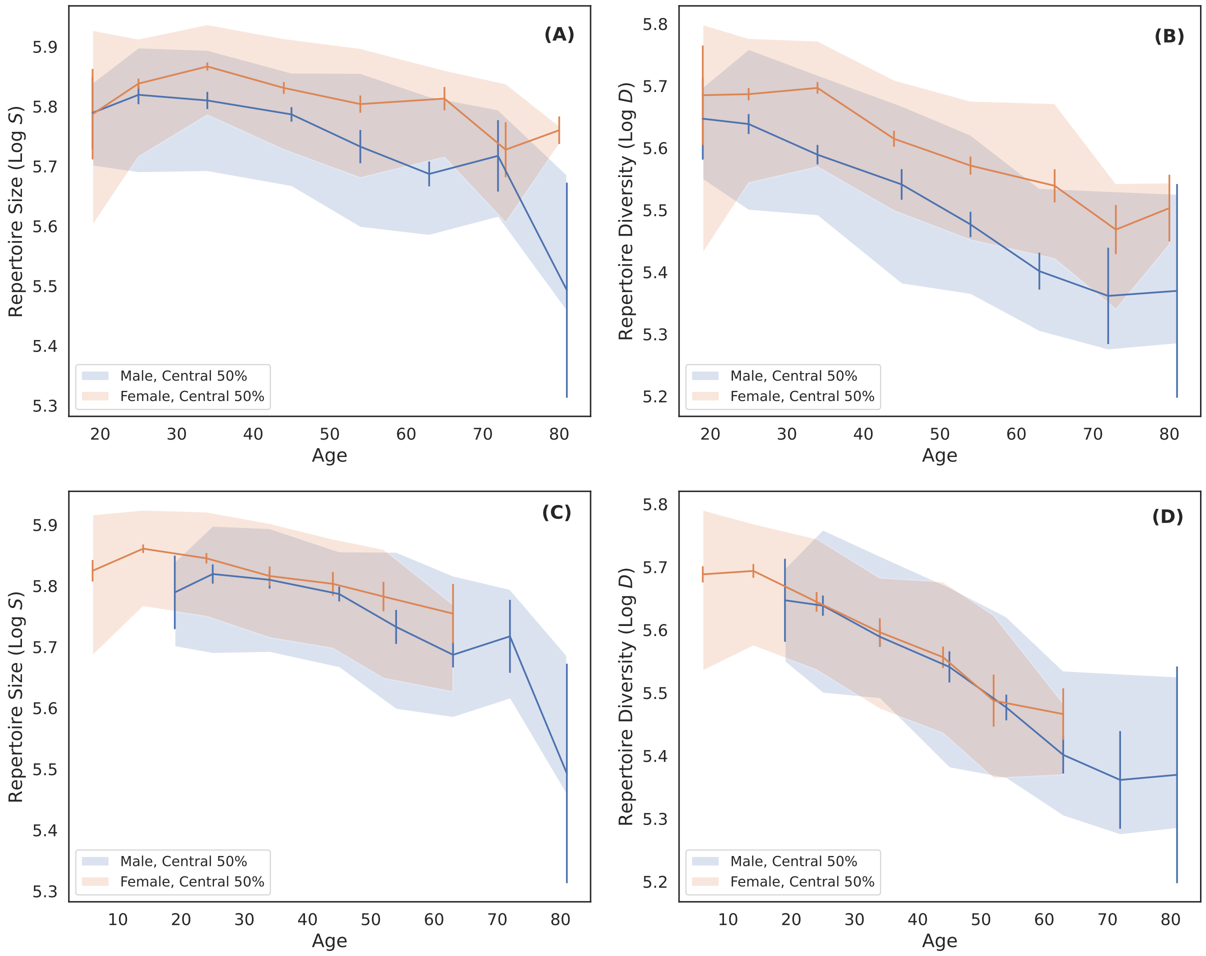


Figure S2: **Repertoire size and TCR diversity as a function of age and sex.** Same as Figure 2 in the main text but using an independent cohort. (A) Log of the total number of productive TCRs sequenced (Log $S$) as a function of age and sex. Blue and orange curves are the median in decade wide age bins for males and females, respectively. Error bars are bootstrapped and shaded regions indicate the distribution of the central 50% of the data points. (B) Log of the TCR diversity as a function of age (Log $D$) for male (blue) and female (orange) subjects. The binning procedure and shaded regions are the same as in (A). (C) and (D) are the same as (A) and (B), respectively, but with female samples shifted to younger ages by 16 years and rebinned for display.

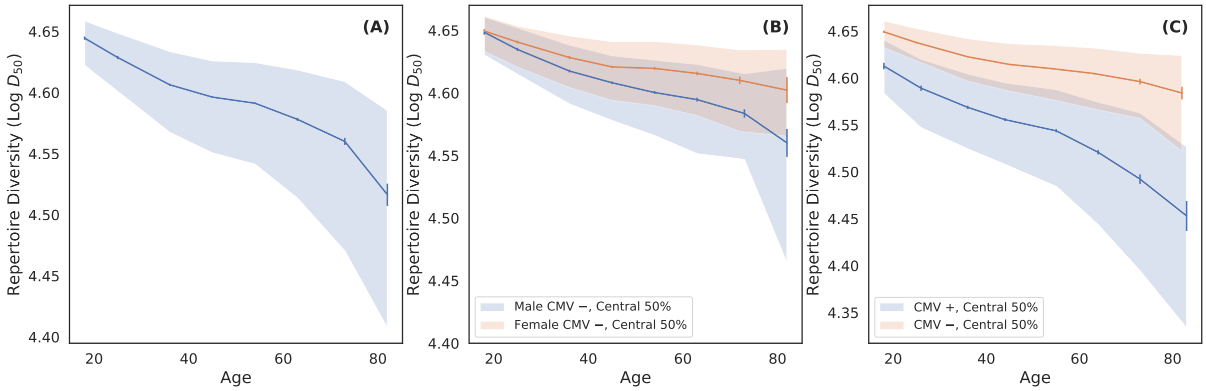


Figure S3: **TCR diversity measured via down-sampling and its dependence on age, sex and CMV status.** (A) Log of the down-sampled TCR diversity (Log $D_{50}$) as a function of age. The blue curve is the median in decade wide age bins, the error bars are bootstrapped and the shaded region indicates the distribution of the central 50% of the data points. (B) Log of the down-sampled TCR diversity as a function of age for male (blue) and female (orange) subjects. All subjects in (B) are CMV negative (see text for details). The binning procedure and shaded regions are the same as in (A). (C) Log of the down-sampled TCR diversity as a function of age for CMV positive (blue) and CMV negative (orange) subjects.

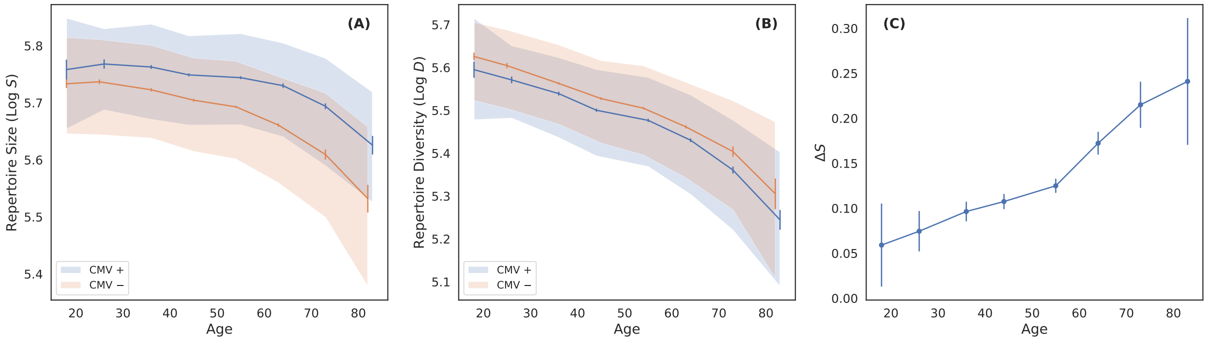


Figure S4: **Impact of CMV on repertoire as a function of age.** (A) Log of the total number of productive TCRs (Log $S$) as a function of age and CMV status. Data are binned in decade age bins. Blue and orange curves are the median in age bins for CMV positive and CMV negative subjects, respectively. Error bars are bootstrapped and shaded regions indicate the distribution of the central 50% of the data points. (B) Log base 10 of the TCR diversity (Log $D$) as a function of age for CMV positive (blue) and CMV negative (orange) subjects. The binning procedure is the same as in (A). (C) Fractional increase in the size of the repertoire due to CMV positivity, i.e. $\Delta S = \frac{S_+ - S_-}{S_-}$. where $S_+$ and $S_-$ are $S$ for CMV postive and CMV negative subjects, respectively. CMV positive subjects have significantly greater number of T cells and the difference increases with age.

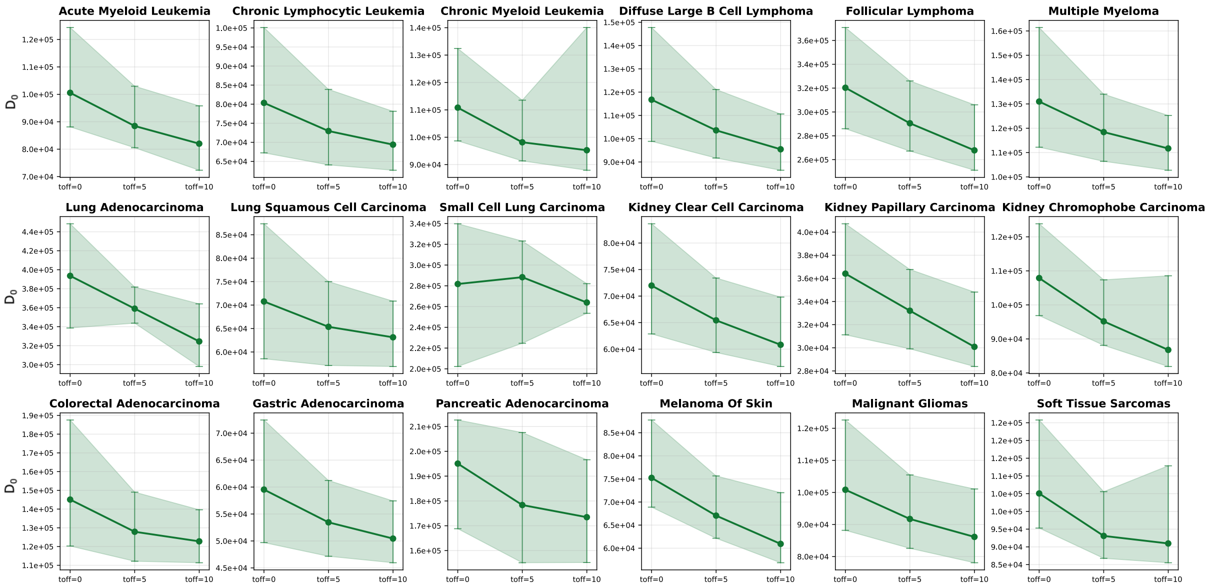


Figure S5: **Covariance between latency offset ( $t_{\text{off}}$) and $D_0$.** Larger values of $toff$ result in smaller values of $D_0$, indicating greater immune contribution to cancer incidence. Across all subtypes, $D_0$ shows minimal systematic variation with $t_{\text{off}}$, demonstrating that covariance effects are smaller than statistical uncertainties.

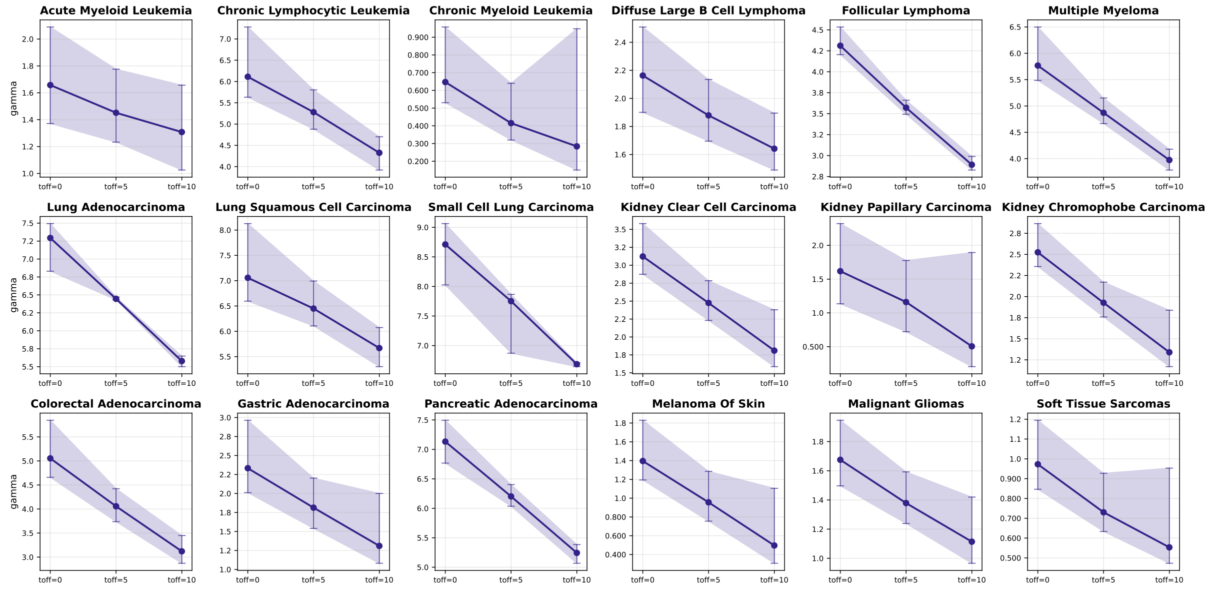


Figure S6: **Covariance between latency offset ( $t_{\text{off}}$) and $\gamma$.** Across subtypes, larger $t_{\text{off}}$ correspond to smaller fitted $\gamma$ values, reflecting covariance between these parameters and a reduced mutational contribution to incidence.

### References


(*1*) Bolen, C. R. et al. (2017). The Repertoire Dissimilarity Index as a method to compare lymphocyte receptor repertoires. *BMC bioinformatics 18*, 1–8.

(*2*) Zuhair, M. et al. (2019). Estimation of the worldwide seroprevalence of cytomegalovirus: a systematic review and meta-analysis. *Reviews in medical virology 29*, e2034.

(*3*) La Rosa, C., and Diamond, D. J. (2012). The immune response to human CMV. *Future virology 7*, 279–293.

(*4*) Khan, N. et al. (2002). Cytomegalovirus seropositivity drives the CD8 T cell repertoire toward greater clonality in healthy elderly individuals. *The Journal of Immunology 169*, 1984–1992.

(*5*) Almanzar, G. et al. (2005). Long-term cytomegalovirus infection leads to significant changes in the composition of the CD8+ T-cell repertoire, which may be the basis for an imbalance in the cytokine production profile in elderly persons. *Journal of virology 79*, 3675–3683.

(*6*) Pourgheysari, B. et al. (2007). The cytomegalovirus-specific CD4+ T-cell response expands with age and markedly alters the CD4+ T-cell repertoire. *Journal of virology 81*, 7759–7765.

(*7*) Sansoni, P. et al. (2014). New advances in CMV and immunosenescence. *Experimental gerontology 55*, 54–62.

(*8*) Furman, D. et al. (2015). Cytomegalovirus infection enhances the immune response to influenza. *Science translational medicine 7*, 281ra43–281ra43.

(*9*) Wang, G. C. et al. (2012). T cell receptor $\alpha\beta$ diversity inversely correlates with pathogen-specific antibody levels in human cytomegalovirus infection. *Science translational medicine 4*, 128ra42–128ra42.

(*10*) Van Leeuwen, E. M. et al. (2006). Differential usage of cellular niches by cytomegalovirus versus EBV-and influenza virus-specific CD8+ T cells. *The Journal of Immunology 177*, 4998–5005.

(*11*) Wertheimer, A. M. et al. (2014). Aging and cytomegalovirus infection differentially and jointly affect distinct circulating T cell subsets in humans. *The Journal of Immunology 192*, 2143–2155.

(*12*) Lindau, P. et al. (2019). Cytomegalovirus exposure in the elderly does not reduce CD8 T cell repertoire diversity. *The Journal of Immunology 202*, 476–483.

(*13*) Emerson, R. O. et al. (2017). Immunosequencing identifies signatures of cytomegalovirus exposure history and HLA-mediated effects on the T cell repertoire. *Nature genetics 49*, 659–665.

(*14*) Greissl, J. et al. (2021). Immunosequencing of the T-cell receptor repertoire reveals signatures specific for diagnosis and characterization of early Lyme disease. *medRxiv*.

(*15*) Elyanow, R. et al. (2022). T cell receptor sequencing identifies prior SARS-CoV-2 infection and correlates with neutralizing antibodies and disease severity. *JCI insight 7*.
